# Infusion of CCR5 Gene-Edited T Cells Allows Immune Reconstitution, HIV Reservoir Decay, and Long-Term Virological Control

**DOI:** 10.1101/2021.02.28.433290

**Authors:** Joumana Zeidan, Ashish A. Sharma, Gary Lee, Angie Raad, Remi Fromentin, Slim Fourati, Khader Ghneim, Gabriela P. Sanchez, Clarisse Benne, Glenda Canderan, Francesco A. Procopio, Robert Balderas, Georges Monette, Jacob P. Lalezari, Jane M. Heffernan, Laurent Sabbagh, Nicolas Chomont, Dale Ando, Steven G. Deeks, Rafick-Pierre Sekaly

## Abstract

Antiretroviral therapy (ART) fails to fully restore immune function and is not curative. A single infusion of CCR5 gene-edited autologous CD4^+^ T cells (SB-728-T) led to sustained increases in CD4^+^ T cell counts, improved T cell homeostasis, and reduced the estimated size of the HIV reservoir. These outcomes were associated with the expansion and long-term persistence of a novel CCR5 gene-edited CD4^+^ T memory stem cell (CD45RA^int^RO^int^ T_SCM_) subset that can replenish the pool of more differentiated memory cells. We showed that novel CD45RA^int^RO^int^ T_SCM_ cells are transcriptionally distinct from the previously described CD45RA^+^ T_SCM_ and are minimally differentiated cells uncommitted to a specific Th-lineage. Subsequently, we showed in an independent trial that infusion of the SB-728-T cell product resulted in partial control of viral replication upon cessation of ART which was correlated with the frequencies of CCR5 gene-edited T_SCM_ and their T_EM_ progeny. Interestingly, one participant that remained off ART to this date demonstrated long-term maintenance of CCR5 gene-edited cells and increased frequency of polyfunctional HIV-specific CD4^+^ and CD8^+^ T cells, contributing to low levels of viral load 5 years post-infusion. Consequently, the generation of HIV protected memory CD4^+^ T cells by CCR5 disruption can contribute toward novel interventions aimed at achieving a sustained ART-free viral remission of HIV disease.

## Introduction

Although ART can durably suppress viral replication, HIV persists indefinitely, requiring infected individuals to remain on complex antiretroviral drug regimens for life. The ability of ART to reconstitute immune function is highly variable. A subset of individuals who initiate ART later in the disease course (10% to 45%, i.e., “immune non-responders”) fail to exhibit complete restoration of CD4^+^ T cell counts even after years of effective ART^1^. Diminished CD4^+^ T cell recovery has been associated with several host-related and HIV-related factors such as impaired thymopoiesis and homeostasis^2–4^. As low CD4^+^ T cell counts in individuals on ART have been associated with increased risk of cardiovascular complications, cancer and other comorbidities^5–8^, novel therapeutic approaches to restore immune homeostasis are needed.

Studies quantifying the latent HIV reservoir have shown minimal decay of total and integrated HIV DNA four years post ART initiation, particularly in participants who received ART only during the chronic phase of infection^9,10^. Several mechanisms contribute to HIV persistence including “latent” infection of long-lived memory CD4^+^ T cells^11–13^ that are maintained by homeostatic proliferation^14–16^ and dysfunctional host clearance mechanisms^2,17^. Intriguingly, these mechanisms are exacerbated in immune non-responders^3,18,19^ and have been associated with higher frequencies of HIV-infected cells^20^. Therefore, enhancing the recovery of CD4^+^ T cells may contribute to the reduction of the HIV reservoir during ART.

CCR5 is one of the major co-receptors for HIV entry. The therapeutic advantage of providing HIV-infected individuals with a CCR5-deficient immune compartment was demonstrated with the “Berlin Patient”^21,22^, who was HIV-free since receiving allogeneic bone marrow transplants of CD34^+^ stem cells from a homozygous CCR5Δ32 matched donor. While these results are encouraging, a less invasive and a more broadly applicable curative strategy would be desirable. One approach is to reconstitute immune function through adoptive transfer of autologous T cells which has shown promising results in other viral infections, including cytomegalovirus and Epstein-Barr virus^23,24^, but largely failed in HIV infection^25–28^, partly because CD4^+^ T cells remain susceptible to HIV infection. Perez *et al.,* demonstrated that HIV-infected NOD/SCID/IL-2Rγ^null^ mice transplanted with CCR5 gene-edited cells had higher CD4^+^ T cells and lower plasma viremia compared to mice that received mock CD4^+^ T cells^29^. In addition, a clinical trial with adoptively transferred zinc finger nuclease (ZFN)-mediated CCR5 gene-edited CD4^+^ T cells (SB-728-T products) in HIV-infected adults demonstrated that infusion was safe, well tolerated and led to increased CD4^+^ T cell counts^30^. Herein, we show in two independent clinical trials that expansion of CD45RA^int^RO^int^ T_SCM_ and resetting of T cell homeostasis is a mechanism that underlies the long-term benefits of this intervention.

## Results

### A novel memory stem cell-like CD4^+^ T cell subset contributes to restoration of T cell homeostasis and correlates with reservoir decay

The clinical study SB-728-0902 evaluated nine HIV-infected immune non-responders on long-term ART (7-22 years) who had at baseline a mean CD4^+^ T cell count of 363 cells/µL (**Supplementary Table 1**).

All participants received a single infusion of 1×10^10^ to 3×10^10^ SB-728-T product containing between 14% to 36% ZFN-mediated CCR5 gene-edited alleles (3 cohorts, *n* = 9; see online Methods and **Supplementary Table 2**). The infused product was devoid of naïve cells and included mostly (34% ± 16) effector memory cells (T_EM_) and a population that expressed intermediate levels of both CD45RA and CD45RO (CD45RA^int^RO^int^; mean of 31% ± 17). A subset of CD45RA^int^RO^int^ cells (14.5% ± 7.05) also expressed CD27 and CCR7 (**Supplementary Fig. 1a, b**). CD45RA^int^RO^int^ CD27^+^CCCR7^+^ cells had the highest levels of gene edited alleles in SB-728-T (45.9% *vs.* 37.9% in T_EM_, *P* = 0.002; **Supplementary Fig. 1c**), as determined by deep sequencing of the CCR5 allele. The frequency of cells harboring integrated HIV DNA in the product was significantly lower than that of pre-manufacture cells (*P* = 0.0039; **Supplementary Fig. 1d**). The frequency of integrated HIV DNA was significantly lower in CD45RA^int^RO^int^ CD27^+^CCR7^+^ cells from SB-728-T products than in other memory subsets (mean of 58.11 copy/10e^6^ cells *vs.* 679.4 copy/10e^6^ cells in T_EM_, *P* = 0.0078; **Supplementary Fig. 1e**).

CCR5 gene-edited cells expanded post infusion and peaked at 7-21 days (median 2.4-fold expansion at 21 days; **Supplementary Fig. 2a**). CCR5 gene-edited CD4^+^ T cells, assessed by the Pentamer Duplication assay (a measure of a specific five-nucleotide insertion at the site of ZFN-mediated editing that represents ∼25% of all modified sequences^29,30^) and DNA-Seq of the CCR5 locus, were detected up to 3-4 years in PBMCs (**Supplementary Table 2**), with frequencies of gene-edited alleles in CD4^+^ T cells ranging between 5 and 15.7% (*n* = 5; **Supplementary Fig. 2b**), and up to 12 months (last sampled time point) in rectal mucosal biopsies (**Supplementary Fig. 2c**). In addition, the frequency of edited mononuclear cells in lymph node tissues (*n* = 3) was similar to that found in the periphery (**Supplementary Fig. 2d**). Consequently, peripheral CD4^+^ T cells counts also increased post-infusion (**Supplementary Table 2 and Supplementary Fig. 2e**), similar to the findings of Tebas *et al.*^30^, and remained significantly above baseline (BL) for 3-4 years post-infusion (+162 cells/µL, *P* = 0.02; **Supplementary Fig. 2e**). The infusion dose (CCR5 gene-edited cell numbers) did not correlate with peak or long-term CD4^+^ T cell counts (*P* = 0.95 and *P* = 0.91; data not shown). Increased CD4^+^ T cell counts and the restoration of the CD4:CD8 ratio (mean of

0.62 at BL *vs.* at 0.84 month 12, *P* = 0.0078; data not shown) were associated with the expansion of gene-edited cells post-infusion (**Supplementary Table 3)**.

To identify the mechanisms leading to the reconstitution of CD4^+^ T cells, a longitudinal analysis of the distribution of CD4^+^ T cell subsets post-infusion was performed. We found a specific increase in the frequency and absolute numbers of CD4^+^ T cells expressing intermediate levels of CD45RA and CD45RO (termed “CD45RA^int^RO^int^”) at every time point analyzed post-infusion (from 13.8% at BL to 38.9% at day 14-28 (*P* = 0.03) and 25.4% at year 3-4 (*P* = 0.008); **Fig. 1a and Supplementary Fig. 3a and b**). CD45RA^int^RO^int^ cells expressed markers previously shown to be up-regulated on CD45RA^+^ memory stem cells (CD45RA^+^ T_SCM_)^31^, such as CD127, CD28, CD58 and CD95^32^ (**Fig. 1b**), suggesting that this subset could represent a novel T_SCM_ subset. Increased CD45RA^int^RO^int^ T_SCM_ counts (**Supplementary Fig. 3c**), but not that of any other memory subsets, were significantly correlated with long-term increases of CD4^+^ T cell counts (**Table 1**) as well as with long-term frequencies of CCR5 gene-edited CD4^+^ T cells (*P* = 0.0002, **Supplementary Fig. 3d**). Consequently, levels of the Pentamer Duplication marker were specifically enriched within CD45RA^int^RO^int^ T_SCM_ post-infusion with a mean of 10- and 38-fold higher levels in CD45RA^int^RO^int^ T_SCM_ compared to central memory (T_CM_) or effector memory (T_EM_) cells at years 3-4, respectively (**Fig. 1c**). Sequencing of CCR5 DNA mutations confirmed the long-term enrichment of CCR5 gene-edited alleles in CD45RA^int^RO^int^ T_SCM_ (23% ± 11 CCR5 gene-edited alleles at year 3-4 compared to 7.9% ± 5.1 in T_CM_ (*P* = 0.02) and 5.9% ± 6.6 in T_TM_ (*P* = 0.02); **Supplementary Fig. 3e**). Importantly, expansion of CD45RA^int^RO^int^ T_SCM_ post infusion was associated with a polyclonal reconstitution of the CD4+ T cell compartment as no changes were observed in CCR5 insertion/deletion (indel) diversity (**Supplementary Fig. 3f**) and in TCR diversity, as measured by similar Shannon entropy indexes^33^ (**Supplementary Fig. 3g**), post infusion.

**Figure 1.**
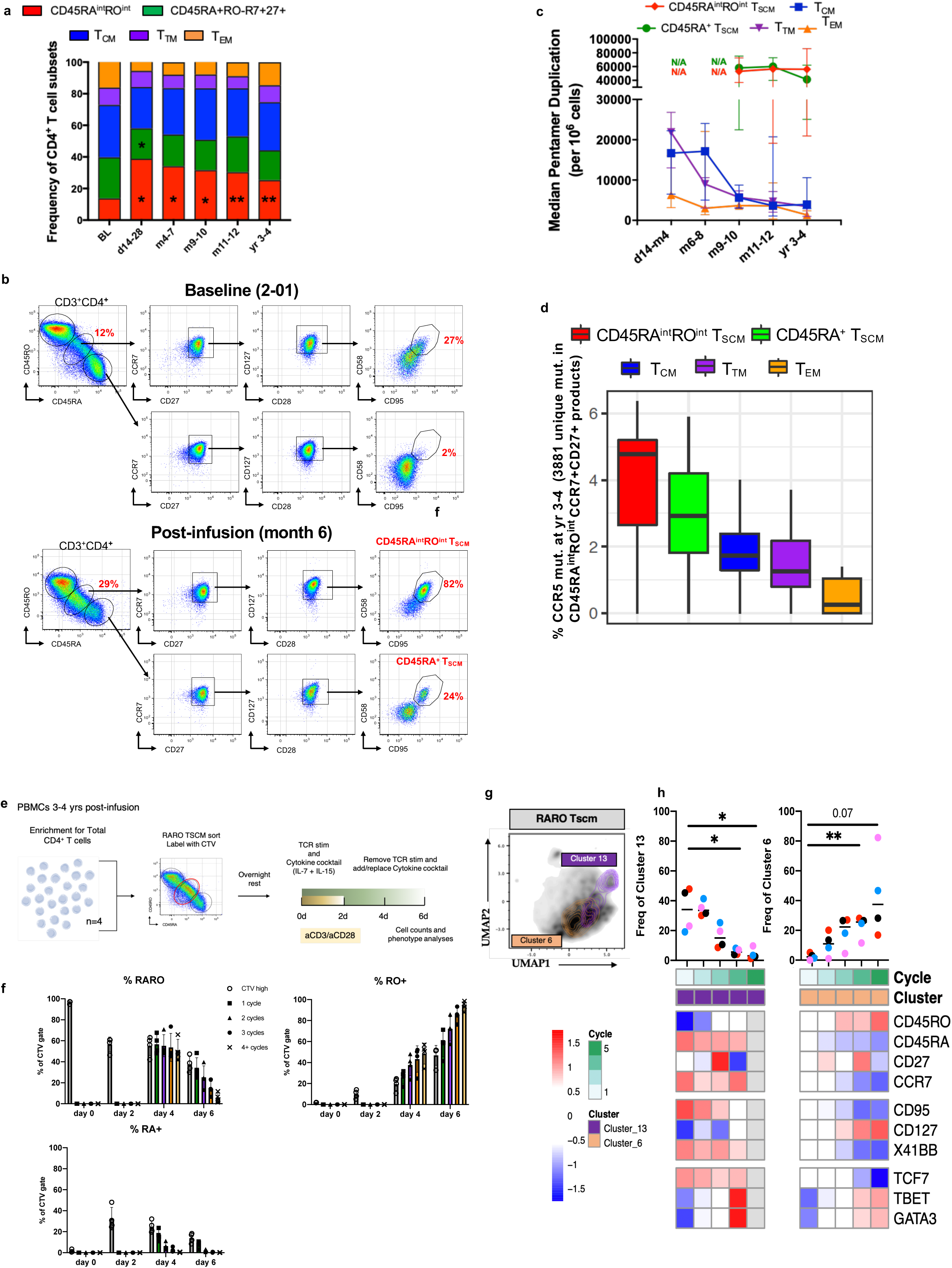
Identification of a novel memory stem cell CD4^+^ T cell subset (CD45RA^int^RO^int^ cells expressing CD95) that contributes to the persistence of CCR5 gene-edited T cells and total CD4^+^ T cells. **a,** Bar chart depicting the mean distribution of CD45RA^+^RO^−^R7^+^27^+^ cells (that include naïve and CD45RA^+^ T_SCM_), T_CM_, T_TM_, T_EM_, and CD45RA^int^RO^int^ frequencies in CD4^+^ T cells at BL (*n* =9), early (days 14-28; *n* = 6), mid (months 4-7, *n* = 7, and months 9-10, *n* = 7), late (months 11-12, *n* = 9), and long-term time points (year 3-4, *n* = 9) post-infusion. * *P* < 0.05, ** *P* < 0.01; Wilcoxon rank-sum test. **b,** Representative example of the gating strategy used to analyze the expression of CD58 and CD95, markers upregulated by memory stem cells, in CD45RA^int^RO^int^ and CD45RA^+^RO^−^ subsets for participants 2-01 at BL and 6 months post SB-728-T infusion. **c,** Median frequency of the Pentamer Duplication marker per 10^6^ cells, a specific sequence tag that accounts for approximately 25% of CCR5 gene-edited cells, measured in sorted T_CM_, T_TM_, and T_EM_ memory subsets at d14-m4 (*n* = 7 for all 3 subsets), m6-8 (*n* = 7, 7, and 6, respectively), m11-12 (*n* = 7, 7, and 6, respectively), and year 3-4 (*n* = 7, 7, and 5, respectively), and in CD45RA^+^ T_SCM_ and CD45RA^int^RO^int^ T_SCM_ at m9-10 (*n* = 6 and 5, respectively), m11-12 (*n* = 3 and 5, respectively), and year 3-4 (*n* = 7 and 8, respectively) post-infusion. N/A = not done due to limitations in availability of cryopreserved PBMCs. **d,** Distribution of the ZFN-mediated CCR5 mutations, determined by DNA sequencing, present uniquely in CD45RA^int^CD45RO^int^ CCR7^+^CD27^+^ cells from SB-728-T products in CD4^+^ T cell subsets at year 3-4 (*n* = 5). Box shows median, first and third quartiles, and whiskers extend to a distance of 1.5*IQR. **e-g,** *In vitro* stimulation of purified CD45RA^int^RO^int^ T_SCM_ sorted from year 3-4 samples (*n* = 4) post-infusion with DynaBeads Human T-activator CD3/CD28. Frequency of CD45RA^int^RO^int^, CD45RA^+^, and CD45RO^+^ cells in culture up to 6 days post stimulation in the different cycles of proliferation (**f**). Uniform Manifold Approximation and Projection (UMAP) dimension reduction analysis of flow cytometric phenotypic analysis of cells emerging from the CD45RA^int^RO^int^ T_SCM_ subset post stimulation, with overlays of clusters 13 (which demonstrates lowest cycling at day 6) and 6 (which demonstrates highest cycling at day 6) (**g**). Frequency of clusters 13 and 6 for each proliferation cycle (1-5) and heatmap of their marker co-expression with red and blue squares indicating upregulation and down-regulation, respectively (**h**). Columns represent different cycles.

**Table 1.**
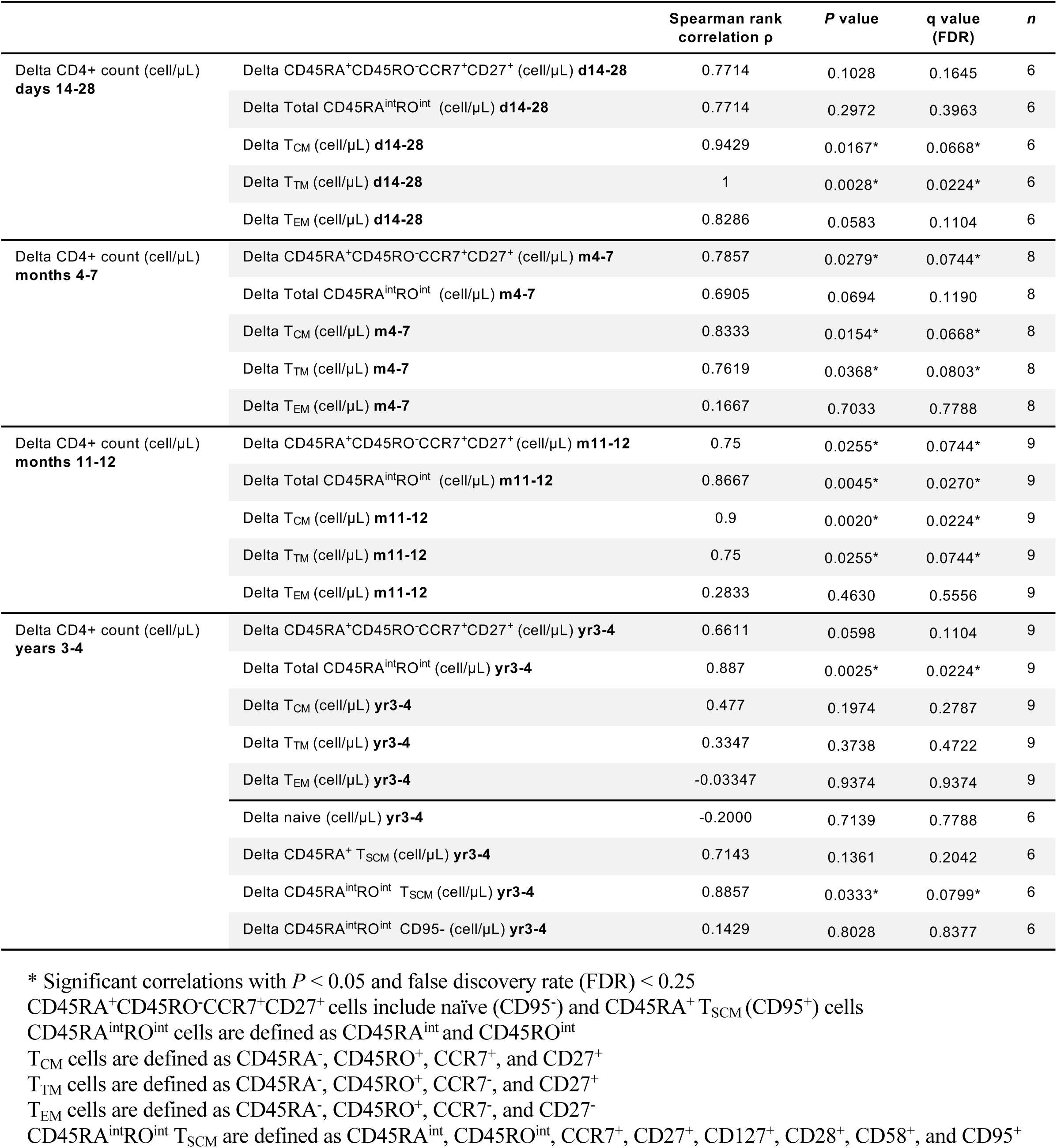
Correlation of increase in circulating CD4^+^ T cell subsets (delta cell count compared to BL) at early (day 14), mid (month 4-7), late (month 11-12) and long-term time points (years 3-4) with immune reconstitution (delta CD4^+^ count compared to BL) at the same time points.

| | | Spearman rank correlation $\rho$ | *P* value | q value (FDR) | *n* |
| --- | --- | --- | --- | --- | --- |
| Delta CD4<sup>+</sup> count (cell/μL)<br>**days 14-28** | Delta CD45RA<sup>+</sup>CD45RO<sup>-</sup>CCR7<sup>+</sup>CD27<sup>+</sup> (cell/μL) **d14-28** | 0.7714 | 0.1028 | 0.1645 | 6 |
| | Delta Total CD45RA<sup>int</sup>RO<sup>int</sup> (cell/μL) **d14-28** | 0.7714 | 0.2972 | 0.3963 | 6 |
| | Delta T<sub>CM</sub> (cell/μL) **d14-28** | 0.9429 | 0.0167* | 0.0668* | 6 |
| | Delta T<sub>TM</sub> (cell/μL) **d14-28** | 1 | 0.0028* | 0.0224* | 6 |
| | Delta T<sub>EM</sub> (cell/μL) **d14-28** | 0.8286 | 0.0583 | 0.1104 | 6 |
| Delta CD4<sup>+</sup> count (cell/μL)<br>**months 4-7** | Delta CD45RA<sup>+</sup>CD45RO<sup>-</sup>CCR7<sup>+</sup>CD27<sup>+</sup> (cell/μL) **m4-7** | 0.7857 | 0.0279* | 0.0744* | 8 |
| | Delta Total CD45RA<sup>int</sup>RO<sup>int</sup> (cell/μL) **m4-7** | 0.6905 | 0.0694 | 0.1190 | 8 |
| | Delta T<sub>CM</sub> (cell/μL) **m4-7** | 0.8333 | 0.0154* | 0.0668* | 8 |
| | Delta T<sub>TM</sub> (cell/μL) **m4-7** | 0.7619 | 0.0368* | 0.0803* | 8 |
| | Delta T<sub>EM</sub> (cell/μL) **m4-7** | 0.1667 | 0.7033 | 0.7788 | 8 |
| Delta CD4<sup>+</sup> count (cell/μL)<br>**months 11-12** | Delta CD45RA<sup>+</sup>CD45RO<sup>-</sup>CCR7<sup>+</sup>CD27<sup>+</sup> (cell/μL) **m11-12** | 0.75 | 0.0255* | 0.0744* | 9 |
| | Delta Total CD45RA<sup>int</sup>RO<sup>int</sup> (cell/μL) **m11-12** | 0.8667 | 0.0045* | 0.0270* | 9 |
| | Delta T<sub>CM</sub> (cell/μL) **m11-12** | 0.9 | 0.0020* | 0.0224* | 9 |
| | Delta T<sub>TM</sub> (cell/μL) **m11-12** | 0.75 | 0.0255* | 0.0744* | 9 |
| | Delta T<sub>EM</sub> (cell/μL) **m11-12** | 0.2833 | 0.4630 | 0.5556 | 9 |
| Delta CD4<sup>+</sup> count (cell/μL)<br>**years 3-4** | Delta CD45RA<sup>+</sup>CD45RO<sup>-</sup>CCR7<sup>+</sup>CD27<sup>+</sup> (cell/μL) **yr3-4** | 0.6611 | 0.0598 | 0.1104 | 9 |
| | Delta Total CD45RA<sup>int</sup>RO<sup>int</sup> (cell/μL) **yr3-4** | 0.887 | 0.0025* | 0.0224* | 9 |
| | Delta T<sub>CM</sub> (cell/μL) **yr3-4** | 0.477 | 0.1974 | 0.2787 | 9 |
| | Delta T<sub>TM</sub> (cell/μL) **yr3-4** | 0.3347 | 0.3738 | 0.4722 | 9 |
| | Delta T<sub>EM</sub> (cell/μL) **yr3-4** | -0.03347 | 0.9374 | 0.9374 | 9 |
| | Delta naive (cell/μL) **yr3-4** | -0.2000 | 0.7139 | 0.7788 | 6 |
| | Delta CD45RA<sup>+</sup> T<sub>SCM</sub> (cell/μL) **yr3-4** | 0.7143 | 0.1361 | 0.2042 | 6 |
| | Delta CD45RA<sup>int</sup>RO<sup>int</sup> T<sub>SCM</sub> (cell/μL) **yr3-4** | 0.8857 | 0.0333* | 0.0799* | 6 |
| | Delta CD45RA<sup>int</sup>RO<sup>int</sup> CD95<sup>-</sup> (cell/μL) **yr3-4** | 0.1429 | 0.8028 | 0.8377 | 6 |

\* Significant correlations with *P* < 0.05 and false discovery rate (FDR) < 0.25
CD45RA<sup>+</sup>CD45RO<sup>-</sup>CCR7<sup>+</sup>CD27<sup>+</sup> cells include naïve (CD95<sup>-</sup>) and CD45RA<sup>+</sup> T<sub>SCM</sub> (CD95<sup>+</sup>) cells
CD45RA<sup>int</sup>RO<sup>int</sup> cells are defined as CD45RA<sup>int</sup> and CD45RO<sup>int</sup>
T<sub>CM</sub> cells are defined as CD45RA<sup>-</sup>, CD45RO<sup>+</sup>, CCR7<sup>+</sup>, and CD27<sup>+</sup>
T<sub>TM</sub> cells are defined as CD45RA<sup>-</sup>, CD45RO<sup>+</sup>, CCR7<sup>-</sup>, and CD27<sup>+</sup>
T<sub>EM</sub> cells are defined as CD45RA<sup>-</sup>, CD45RO<sup>+</sup>, CCR7<sup>-</sup>, and CD27<sup>-</sup>
CD45RA<sup>int</sup>RO<sup>int</sup> T<sub>SCM</sub> are defined as CD45RA<sup>int</sup>, CD45RO<sup>int</sup>, CCR7<sup>+</sup>, CD27<sup>+</sup>, CD127<sup>+</sup>, CD28<sup>+</sup>, CD58<sup>+</sup>, and CD95<sup>+</sup>

The presence of CCR5 gene mutations in short-lived memory cells such as T_EM_ at years 3-4 post-infusion (0.7% to 3% CCR5 gene-edited alleles; **Supplementary Fig. 3e**) suggested that CCR5 gene-edited cells within long-lived memory cells such as T_SCM_ can differentiate and maintain a small subset of CCR5 gene-edited T_EM_ cells years after the initial infusion. We identified CCR5 ZFN-mediated indels unique to the CD45RA^int^RO^int^ T_SCM_ subset in SB-728-T products (*n* = 3,881) that remained detected in CD45RA^int^RO^int^ T_SCM_ at 3-4 years post infusion, demonstrating their capacity to persist long-term. Additionally, CD45RA^int^RO^int^ T_SCM_-unique indels were detected in other memory T cell subsets post-infusion including short-lived T_EM_ at 3-4 years post infusion (0.49% (95% CI: 0%-1.36%); **Fig. 1d**), suggesting the potential of these cells to generate more differentiated cells. The detection of CD45RA^int^RO^int^ T_SCM_-unique indels in the CD45RA^+^ T_SCM_ subset (**Fig. 1d**) as well as the detection of high levels of the Pentamer Duplication marker in these cells (**Fig. 1d**) suggest that the increase in CD45RA^+^ T_SCM_ (**Supplementary Fig. 3c**) post infusion is a result of their differentiation from CD45RA^int^RO^int^ T_SCM_ post infusion and not from homeostatic proliferation.

To investigate the potential of CD45RA^int^RO^int^ T_SCM_ cells to undergo homeostatic expansion or self-renewal and differentiate into other memory subsets following *in vitro* stimulation, CD45RA^int^RO^int^ T_SCM_ cells were sorted at year 3-4, labelled with CellTrace Violet (CTV), and cocultured with anti-CD3/CD28 Dynabeads and homeostatic cytokines (**Fig. 1e**). At days 4 and 6, the majority of cells had undergone proliferation (91% and 97% CTV low cells, respectively; **Supplementary Fig. 4a**). Cells of the CD45RA^int^RO^int^ phenotype remained detected throughout culture while CD45RO^+^ memory T cells increased progressively (**Fig. 1f**). Interestingly, CD45RA^+^ T_SCM_ peaked at day 2 and were detected in cells undergoing 0-1 rounds of proliferation (**Fig. 1f and Supplementary Fig. 4b**), suggesting that this subset has limited self-renewal capacity and/or is more prone to undergo differentiation with increased cell proliferation. Uniform Manifold Approximation and Projection (UMAP), a non-linear dimensionality reduction technique^34^, was employed to investigate the modulation of markers associated with self-renewal and Th-lineage commitment throughout cell division in culture. Two PhenoGraph clusters were identified (**Fig. 1g and Supplementary Fig. 4c**); cluster 13, which was the most abundant cluster in cells with one cycle of proliferation, decreased in frequencies during proliferation and maintained the CD45RA^int^RO^int^ T_SCM_ phenotype and expression of CD27, CCR7, 41BB, and TCF-7, and cluster 6 which increased during proliferation, consisted of the main cluster in cells with 4+ cycles of proliferation, and up-regulated expression of CD45RO, T-bet and GATA-3 (**Fig. 1h and Supplementary Fig. 4d**). All together, these results indicate that CD45RA^int^RO^int^ T_SCM_ cells most likely represent a novel long-lived memory subset that contributes to the long-term polyclonal persistence of CCR5 gene-edited cells and confirm that CD45RA^int^RO^int^ T_SCM_ cells can differentiate into and replenish the pool of more differentiated memory cells.

### A single SB-728-T infusion led to a continuous decrease in the frequency of HIV-infected cells that correlates with the persistence and differentiation of CCR5 gene-edited CD45RA^int^RO^int^ T_SCM_ cells

Infusion of SB-728-T led to significantly lower frequencies of total HIV DNA in PBMCs at 2 years (*P =* 0.02; **Fig. 2a and Supplementary Table 4**) compared to baseline. Moreover, the frequencies of CD4^+^ T cells with integrated HIV DNA significantly declined at 3-4 years post-infusion (*P =* 0.004; **Fig. 2b**) and correlated with the frequencies of total HIV proviruses, as measured by the intact proviral DNA assay (IPDA) (*P =* 0.02; **Fig. 2c**). These results suggest that expansion and persistence of infused CCR5 gene-edited CD4^+^ T cells have an impact on the decay of the HIV reservoir size.

**Figure 2.**
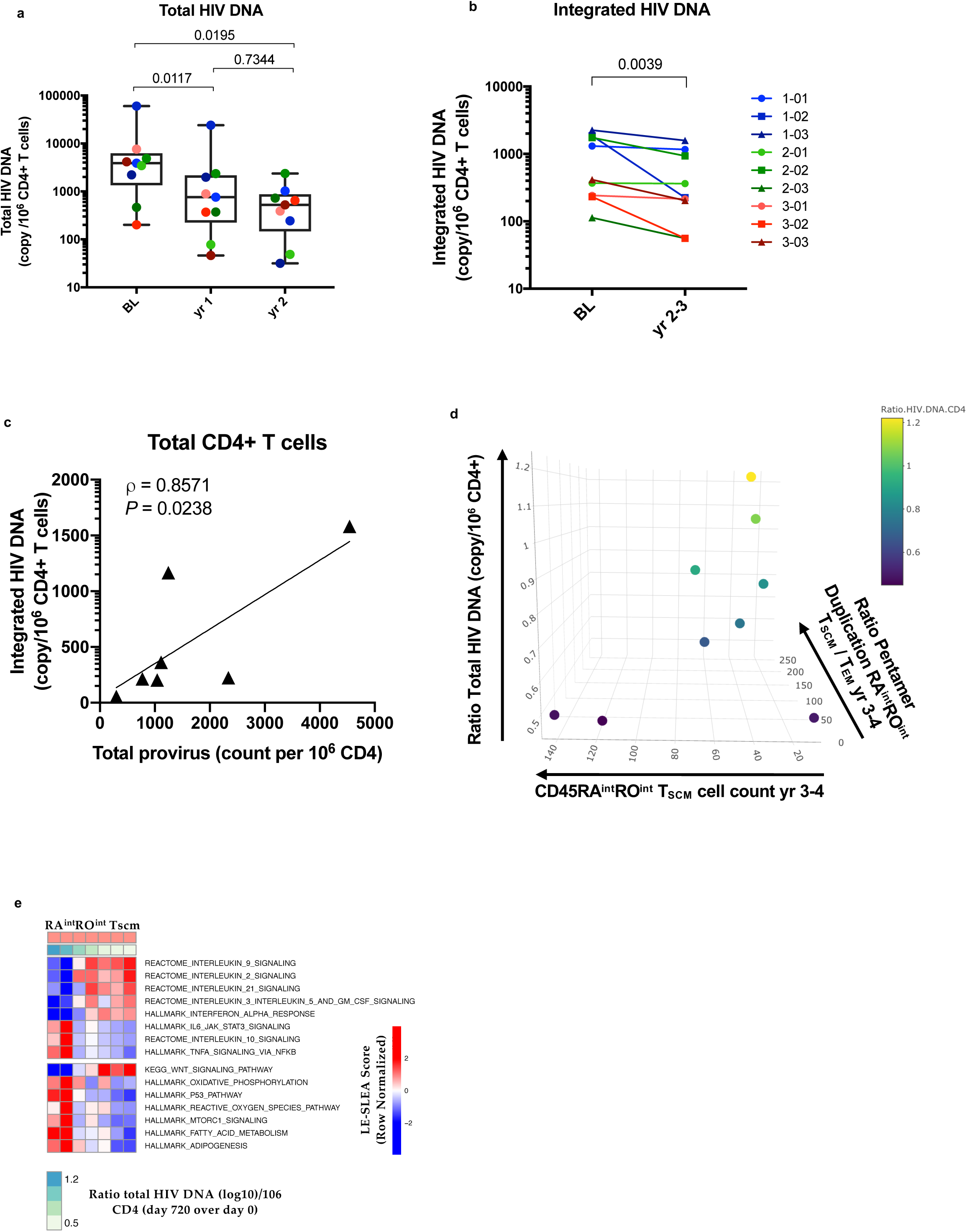
Decay of the HIV reservoir post SB-728-T infusion correlates with persistence of CCR5 gene-edited cells. **a,** Box-plot showing the frequency of cells harboring total HIV DNA per 10^6^ PBMCs at baseline (BL), year 1, and year 2 post-infusion (mean decay of −0.6 log_10_, 95% confidence interval (CI) −1.19 to −0.006 at year 1 and −0.91 log_10_, 95% confidence interval (CI) −1.71 to −0.11 at year 2). Box shows median, first and third quartiles, and whiskers extend to maximum and minimum values. Individual data points are shown for all 9 participants with colors corresponding to the different cohorts (cohort 1, 2 and 3 are shown in blue, green and red hues, respectively). BL values for participants 1-01 and 1-02 were imputed as described in the online Methods. * *P* < 0.05; Wilcoxon rank-sum test. **b,** Frequencies of integrated HIV DNA copies per 10^6^ purified CD4^+^ T cells are shown at BL and year 2-3 (long-term follow up). Participants in cohorts 1, 2 and 3 are shown in blue, green and red symbols, respectively. * *P* < 0.05; Wilcoxon rank-sum test. **c,** Correlation between the frequencies of integrated HIV DNA in CD4^+^ T cells and total HIV provirus assessed by the IDPA assay. **d,** 3-D scatter plot showing the change in the frequency of PBMCs harboring total HIV DNA post-infusion (Ratio of log_10_ values at day 720 over day 0) as a function of CD45RA^int^RO^int^ T_SCM_ cell counts at years 3-4 (z-axis), log_10_ Pentamer Duplication levels in CD45RA^int^RO^int^ T_SCM_ at years 3-4 (x-axis), and the ratio of the frequency of Pentamer Duplication in CD45RA^int^RO^int^ T_SCM_ by the frequency of Pentamer Duplication in T_EM_ at years 3-4 (y-axis). These features, together with log_10_ Pentamer Duplication levels in CD45RA^int^RO^int^ T_SCM_ at year 1 (not plotted), were determined as the best predictors of reservoir decay by a sparse linear multivariate model. The multivariate regression model to predict the reservoir decay contained the following features (see Supplementary Table 5): CD45RA^int^RO^int^ T_SCM_ cell counts at years 3-4, the frequency of Pentamer Duplication in CD45RA^int^RO^int^ T_SCM_ at multiple time points, the Ratio Pentamer Duplication CD45RA^int^RO^int^ T_SCM_/T_EM_ at month 12 and years 3-4, the number of shared mutations between CD45RA^int^RO^int^ T_SCM_ and T_EM_ at years 3-4. Each dot in the scatter plot corresponds to a participant, with dot size proportional to the HIV DNA day 720/BL ratio, with a greater decay (i.e., smaller ratio) symbolized by a greater dot size (adjusted r^2^ = 0.99, F-test: *P* = 0.0008). **e,** Heatmap representing pathways expressed in CD45RA^int^RO^int^ T_SCM_ that were associated with HIV reservoir decay (ratio of log_10_ HIV DNA values at day 720 over day 0; *P* = 0.19), with the range of the outcome presented as legend below the heatmap. Rows represent pathways and columns represent samples. Red and blue correspond to up- and down-regulated pathways respectively. The range of HIV reservoir decay is presented as legends at the bottom of the heatmap.

As CD45RA^int^RO^int^ T_SCM_ cells had significantly lower levels of integrated HIV DNA post-infusion compared to other memory subsets (*P* < 0.05; **Supplementary Fig. 5a**) and minimal contribution to the pool of HIV-infected CD4^+^ T cells (*P* < 0.05; **Supplementary Fig. 5b**) in year 3-4 samples, we investigated the impact of persistence and differentiation of CD45RA^int^RO^int^ T_SCM_ post-infusion on the reservoir decay, using a sparse linear multivariate model to identify features (**Supplementary Table 5**) that could predict the change in the frequency of PBMCs harboring total HIV DNA post-infusion.

This model indicated that a greater decay in the HIV reservoir post-infusion was best predicted by higher CD45RA^int^RO^int^ T_SCM_ cell counts at years 3-4 (*P* = 0.002), higher frequencies of Pentamer Duplication in CD45RA^int^RO^int^ T_SCM_ at years 3-4 (*P* = 0.005), and a lower ratio of the frequency of Pentamer Duplication in CD45RA^int^RO^int^ T_SCM_ over the frequency of Pentamer Duplication in T_EM_ at years 3-4 (*P* = 0.001), reflecting the differentiation of CCR5 gene-edited CD45RA^int^RO^int^ T_SCM_ into the T_EM_ subset (adjusted r^2^ = 0.99, F-test: *P* = 0.0008; **Fig. 2d**).

Linear regression analysis of genesets of sorted CD45RA^int^RO^int^ T_SCM_ cells post infusion with the decay of HIV reservoir (total HIV DNA ratio) showed that pathways associated with homeostatic proliferation (STAT-5 signaling) and stemness (Wnt signaling) were correlated with a greater decay in HIV DNA 2 years post infusion (**Fig. 2e**), demonstrating that the pathways crucial for T_SCM_ homeostasis are also involved in the observed decay of HIV-infected cells^35,36^ (**Supplementary Fig. 5c and Supplementary Table 6**). As CD45RA^int^RO^int^ T_SCM_ cells harbor significantly lower levels of HIV-infected cells than other memory subsets (**Supplementary Figs. 1e, 5a, 5b**), our results suggest that long-term persistence of CCR5 gene-edited CD45RA^int^RO^int^ T_SCM_ can lead to a reduction in the size of the HIV reservoir as a result of these cells differentiating into and replenishing the pool of more differentiated T_EM_.

### CD45RA^int^RO^int^ T_SCM_ are distinct from the previously described CD45RA^+^ T_SCM_ and are minimally differentiated cells uncommitted to a specific Th-lineage

Transcriptional analysis of sorted CD4^+^ T cell subsets was performed on samples from 3-4 years post-infusion. Multi-dimensional scaling of gene expression variance and differential gene expression analysis showed greater dissimilarity between CD45RA^int^RO^int^ T_SCM_ and T_CM_ and T_EM_ than with CD45RA^+^ T_SCM_ (**Supplementary Fig. 6a,b**). Our results also indicated that CD45RA^int^RO^int^ T_SCM_ were transcriptionally distinct from the CD45RA^+^ T_SCM_ subset previously described^31,32^. Figure 3a shows that CD45RA^+^ T_SCM_ expressed several genes associated with T cell activation (e.g., TOX, LAG-3) and effector function (e.g., Perforin, IFN-g, Granzyme B). CD45RA^int^RO^int^ T_SCM_ showed upregulated levels of ID3, CCR7, and CD27, all markers of undifferentiated mature memory CD4^+^ T cells^37,38^ (**Fig. 3a**). SLEA analysis (**Fig. 3b** and **Supplementary Table 7**) further demonstrates that CD45RA^int^RO^int^ T_SCM_ show downregulation of genes associated with activation and cell cycling pathways. Definite demonstration that these two subsets are distinct is also shown in Fig. 3b, where the pathways associated with T cell stemness^31^ were found to be enriched in CD45RA^int^RO^int^ T_SCM_ compared to CD45RA^+^ T_SCM_.

**Figure 3.**
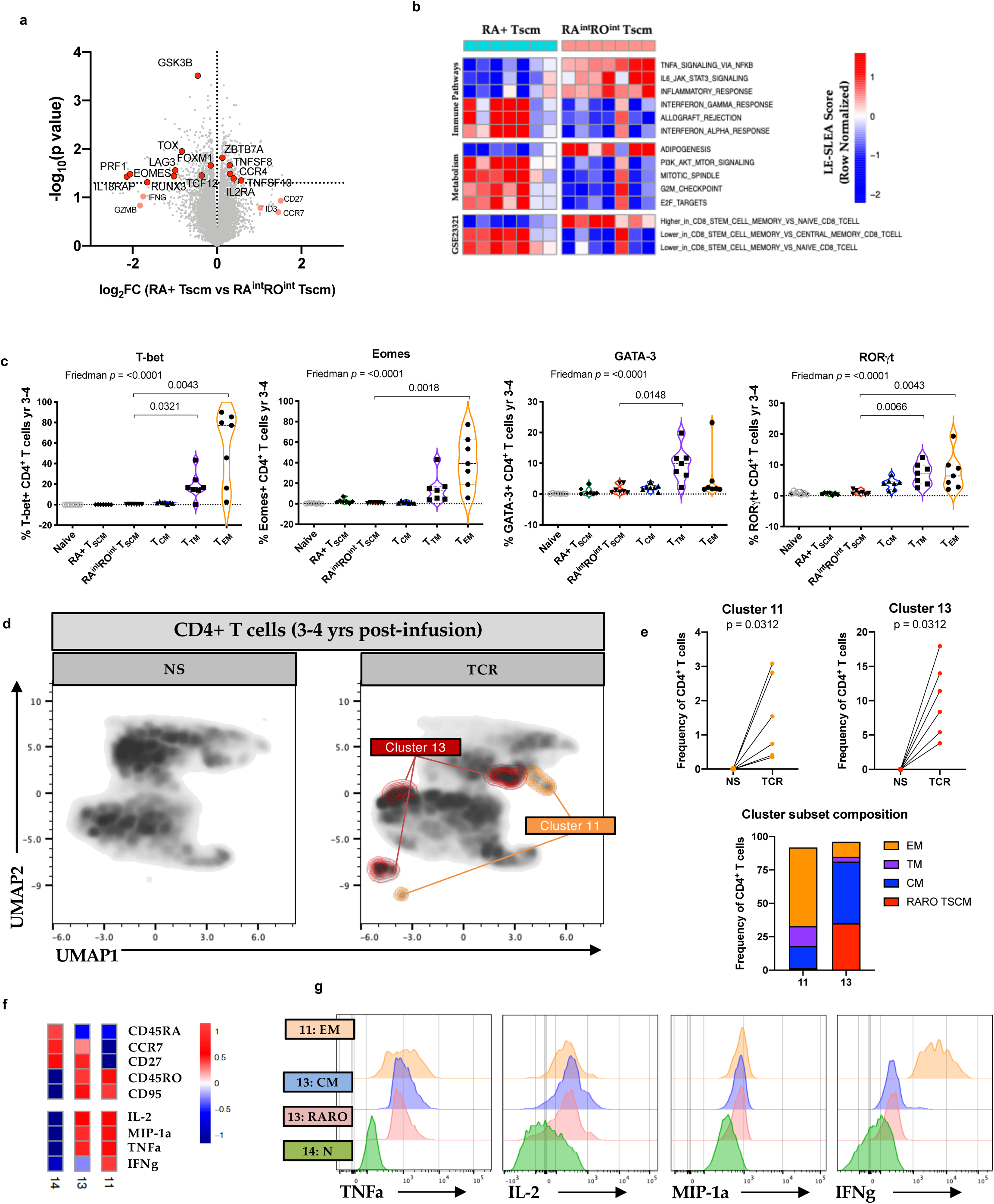
CD45RA^int^RO^int^ T_SCM_ are distinct from previously identified CD45RA^+^ T_SCM_ cells and constitute a novel T_SCM_ subset with features of quiescent uncommitted and long-lived memory cells. **a,** Volcano plot of genes differentially expressed by CD45RA^+^ T_SCM_ and CD45RA^int^RO^int^ T_SCM_ at year 3-4 (*n* = 7) post-infusion. **b,** SLEA plot of selected pathways significantly enriched in genes induced or repressed in CD45RA^int^RO^int^ T_SCM_ compared to CD45RA^+^ T_SCM_ at year 3-4 (*n* = 7) post-infusion. Scale represents SLEA score with red and blue squares indicating positive and negative enrichment respectively. Columns represent CD45RA^int^RO^int^ T_SCM_ and CD45RA^+^ T_SCM_ subsets. **c,** Volcano plots illustrating the expression of the transcription factors T-bet, Eomes, RORγt, and GATA-3 in CD4^+^ T cell subsets at year 3-4 post-infusion (*n* = 7). * *P* < 0.05; Wilcoxon rank-sum test. **d,** Uniform Manifold Approximation and Projection (UMAP) dimension reduction analysis of flow cytometric phenotypic analysis of PBMCs at years 3-4 post infusion (*n = 6*) in response to stimulation with DynaBeads Human T-activator CD3/CD28. **e,** Frequency of two clusters (11 and 13) uniquely up-regulated post-stimulation and their subset distribution. **f,** Heatmap depicting the expression of markers associated with memory T cell phenotypes (CD45RA, CCR7, CD27, CD45RO, and CD95) and effector cytokines (IFN-γ, IL-2, MIP-1α, and TNF-α) of clusters 11 and 13, with a naïve T cell cluster (14) used as control. **g,** MFI levels of TNF-α, IL-2, MIP-1α, and IFN-γ shown for naïve T cells (cluster 14), T_CM_, and CD45RA^int^RO^int^ T_SCM_ (cluster 13) and T_EM_ (cluster 11).

Further indication that CD45RA^int^RO^int^ T_SCM_ cells are uncommitted to a specific Th lineage is demonstrated by the lack of expression of the transcription factors associated with Th1 (T-bet and Eomes), Th2 (GATA-3), and Th17 (RORgt) subsets (**Fig. 3c**). Moreover, upon TCR stimulation of PBMCs at 3-4 years post-infusion, a cluster of cells comprised of CD45RA^int^RO^int^ T_SCM_ produced IL-2, MIP-1a, and TNF-α but not the effector cytokine IFN-γ (**Fig. 3d-g**). In contrast, T_EM_ produced the highest levels of the effector cytokines TNF-α and IFN-γ upon stimulus (**Fig. 3f,g**). Altogether, these results confirmed that CD45RA^int^RO^int^ T_SCM_ cells constitute a novel T_SCM_ subset with features of quiescent uncommitted and long-lived memory CD4^+^ T cells; importantly, these features are associated with the above observed decay of the HIV reservoir as well with immune reconstitution.

### CCR5 gene-edited CD45RA^int^RO^int^ T_SCM_ correlate with control of viral load in participants who underwent treatment interruption 6 weeks post-SB-728-T infusion

We assessed the impact of infusion of CCR5 gene-edited CD4^+^ T cells, including CD45RA^int^RO^int^ T_SCM_, on control of viremia upon cessation of ART in an independent clinical trial (SB-728-1101 study). Participants received escalating doses of cyclophosphamide (CTX) two days prior to infusion (5 cohorts, *n* = 15; see online Methods and **Supplementary Table 8)** and underwent an analytical treatment interruption (ATI) six weeks post-infusion of SB-728-T products. Enrichment of CCR5 mutations within the CD45RA^int^RO^int^ T_SCM_ (27.8% ± 10.4 gene-edited alleles compared to 13.3% ± 4.7 in T_CM_ (*P* = 0.02) and 18.8% ± 6.3 in T_TM_ (*P* = 0.02); **Supplementary Fig. 7a**) was confirmed in SB-728-T products, likely a consequence of their capacity to self-renew and persist. Analysis of viral load (VL) levels showed significantly lower VL during ATI (at week 22) than the historic pre-ART VL set-point (*P =* 0.0054; **Fig. 4a**), indicating that infusion of SB-728-T products may have led to transient but incomplete control of viremia in the majority of the participants. Six individuals who at week 22 showed VL measurements below 10,000 copies/mL and CD4^+^ T cell counts above 500 cells/µl opted to extend ATI beyond week 22 (and remained on ATI for 0.5 – 5 years; **Supplementary Fig. 7b and Supplementary Table 9**). Amongst them, participants who showed a reduction of week 22 VL compared to the historic pre-ART VL greater than 0.5 log were labelled as post-treatment virologic controllers (PTCs). PTCs showed significantly lower levels of infected cells after 16 weeks of ATI than non-controllers, as measured by integrated HIV DNA, as well as total and intact proviruses quantified with the IPDA assay (**Fig. 4b**). Frequencies of CD4^+^ T cells with intact proviruses after 16 weeks of ATI was significantly correlated with the week 22 VL (*P =* 0.0246; **Supplementary Fig. 7c**), indicating that CD4^+^ T cells harboring intact proviruses were contributing to the rebound in VL post ATI.

**Figure 4.**
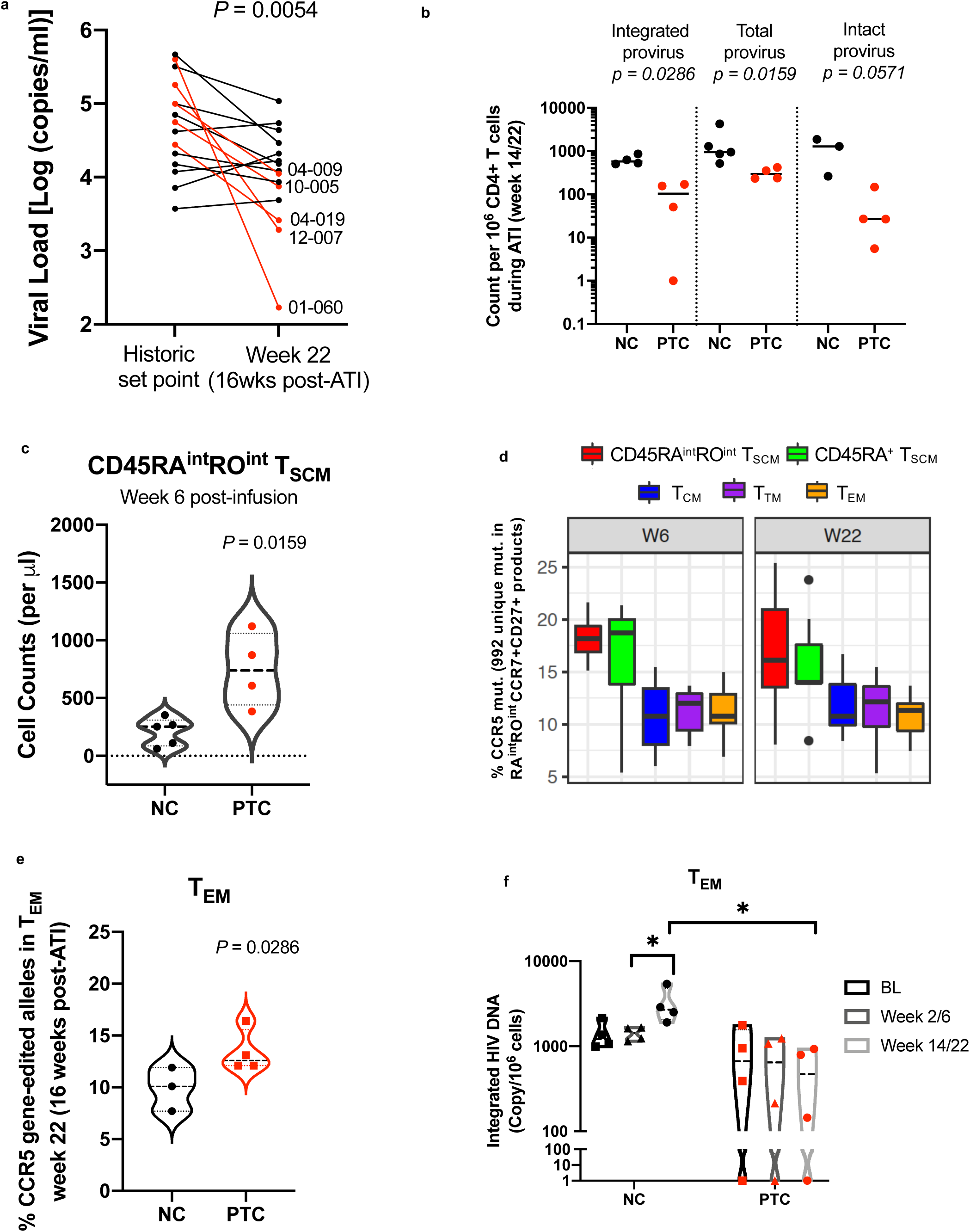
CCR5 gene-edited CD45RA^int^RO^int^ T_SCM_ prior to ATI, levels of CCR5 gene-edited T_EM_ and HIV-specific CD8+ T cell polyfunctionality during ATI correlate with control of viral load and lower reseeding of the T_EM_ HIV reservoir. **a,** Plot depicting the viral load (VL) values at week 22 (equivalent to 16 weeks of ATI) and the historic pre-ART viral set point values obtained from participants’ charts. Participants with extended ATI and a reduction of week 22 VL compared to the historic pre-ART VL greater than 0.5 log are shown in red (post-treatment virologic controllers; PTCs). *P* value of Wilcoxon rank-sum test is shown. VL for all participants post-ATI are shown in Supplementary Table 9. **b,** The HIV reservoir quantified in purified CD4^+^ T cells (integrated HIV DNA, total and intact proviral HIV DNA) after 16 weeks of ATI is shown for non-controllers (black) and PTCs (red). Lines depict means. Virological and immunological assays were performed for participants of cohort 3-5 for whom cryopreserved cells were available for analysis. **c,** Violin plot representing the frequency of CD45RA^int^RO^int^ T_SCM_ cell counts prior to ATI (week 6) in non-controllers (black) and PTCs (red). **d,** Box-plot showing the percent of ZFN-induced CCR5 mutations present uniquely in CD45RA^int^CD45RO^int^ T_SCM_ in SB-728-T products that are detected in CD4^+^ T cell subsets at weeks 6 and 22 post-infusion (*n* = 7). Box shows median, first and third quartiles, and whiskers extend to a distance of 1.5*IQR. Outliers are shown as dots. **e,** Violin plot representing the frequency of CCR5 gene edited alleles in T_EM_ at week 22 in non-controllers (black) and PTCs (red) in participants of cohort 3-5. **f,** Violin plot representing the frequency of integrated HIV DNA within T_EM_ cells at BL, week 6 and week 22 post-infusion (*n* = 7) in non-controllers (black) and PTCs (red) in participants of cohort 3-5. * *P* < 0.05.

Furthermore, PTCs demonstrated significantly higher CD45RA^int^RO^int^ T_SCM_ cell counts prior to ATI (week 6) compared to non-controllers (*P =* 0.016; **Fig. 4c**), indicating a role for these cells in control of VL. To provide an independent quantitative and qualitative assessment of the capacity of CCR5 gene-edited CD45RA^int^RO^int^ T_SCM_ to differentiate into effector memory subsets, we tracked the persistence of CCR5 mutations unique to CD45RA^int^RO^int^ T_SCM_ products in other memory cells post-infusion and post-ATI. We show that CD45RA^int^RO^int^ T_SCM_-unique CCR5 mutations were detected in other CD4^+^ memory subsets at week 6 (10.5%, 11% and 10.9% in T_CM_, in T_TM_ and in T_EM_, respectively) and were maintained at similar frequencies at week 22 (11.7%, 11.3% and 10.7% in T_CM_, in T_TM_ and in T_EM_, respectively; **Fig. 4d**), demonstrating the multipotency capacity of this subset. As T_EM_ cells have been shown to express the highest levels of CCR5 compared to other memory cells^39^, maintenance of a subset of CCR5 gene-edited cells within the T_EM_ subset could lead to protection from *de novo* infection during ATI. To investigate this, we quantified the levels of CCR5 gene edited alleles in CD4^+^ T cell subsets during ATI and found that PTCs showed significantly higher frequencies of CCR5 gene-edited alleles at week 22 exclusively in the T_EM_ subset compared to non-controllers (*P* = 0.029; **Fig. 4e**). Moreover, we measured integrated HIV DNA levels in sorted CD4^+^ T cell subsets at baseline, and at weeks 6 and 22 post-infusion. Our results indicate that the frequency of T_EM_ bearing integrated HIV DNA was significantly increased in non-controllers during ATI (*P* = 0.0083; **Fig. 4f**) but not in PTCs, with PTCs demonstrating significantly lower frequencies of infected T_EM_ cells at week 22 than non-controllers (*P* = 0.0005; **Fig. 4f**). Collectively, these results indicate that continuous replenishment of CCR5 gene-edited T_EM_ cells downstream of differentiation from their CD45RA^int^RO^int^ T_SCM_ precursors confers greater protection from *de novo* infection of the T_EM_ subset in PTCs, consequently having an impact on control of active viral replication during ATI.

### Long-term control of VL in a post-treatment controller that remained off ART for ∼5 years post-infusion is associated with enhanced HIV-specific CD4^+^ and CD8^+^ T cell responses

One of the SB-728-1101 trial participants (01-060) has remained off ART to this date since ART interruption at week 6 post-infusion (∼5 years post infusion). This individual expressed the protective HLA-B57^40–42^ allele and was heterozygote for the delta-32 CCR5 mutation (**Supplementary Table 8**), which may have contributed to his observed VL control. This individual has since been enrolled in the SCOPE cohort (study NCT00187512, conducted at the University of California, San Francisco) and has demonstrated control of VL at low but detectable levels (∼100 copies/mL; **Fig.5a**). CD4^+^ T cell counts have decreased between the last time point (month 12) in the SB-728-1101 trial and his enrollment in the SCOPE cohort to ∼200 cells/µl (**Fig. 5b**); however, this individual declined ART resumption, providing us with a unique opportunity to investigate the role of CCR5 gene-edited cells in partial VL control. Using the Pentamer Duplication assay, we observed higher levels of CCR5 gene-edited cells at 4 years post-infusion compared to month 12 (∼12% of CD4+ cells at year 4 compared to ∼8% at year 1; **Fig 5c**), demonstrating the long-term maintenance of these cells as well as their enrichment compared to the non-edited cells in the presence of prolonged active virus replication. The frequency of CD4^+^ CD45RA^int^RO^int^ T_SCM_ cells remained high post infusion up to month 12 (48% at week 6 and 38% at month 12; **Supplementary Fig. 8a,b**). However, the frequencies of CD4^+^ CD45RA^int^RO^int^ T_SCM_ and T_CM_ progressively decreased post-ATI (from 40% at year 1 to 6.5% at year 4 and from 10.5% at year 1 to 3.6% at year 4, respectively) while that of CD4^+^ T_EM_ and CD8^+^ T_EM_ and T_EMRA_ progressively increased (**Supplementary Fig. 8b**), indicating a pull for differentiation into the T_EM_ and T_EMRA_ subsets in the presence of viremia. To further characterize the heterogeneity of the CD45RA^int^RO^int^ subset, a 22-color Symphony stemness panel was performed on longitudinal samples from patient 01-60 including years 4 and 5 samples. Phenograph analysis identified 2 clusters that increased at early time points post infusion that remained detected long term (clusters 2 and 6, as well as a cluster that was uniquely up-regulated at years 4 and 5 (cluster 13; **Fig. 5d, e and Supplementary Fig. 8c**). Compared to the year 4/5 cluster, the persistent clusters expressed high levels of CCR7, CD27. CD95, CD127, TCF-7, and 4-1BB (**Fig. 5f and Supplementary Fig. 8d,e**), indicating that a subset of cells of the CD45RA^int^RO^int^ phenotype represent long-lasting memory stem cells that can maintain expression of stemness markers during ongoing viral replication. Cells expressing CD45RA^int^RO^int^ but down-regulating CCR7 and CD27 could represent cells transitioning to the T_EM_ phenotype during ongoing viral replication. To determine the impact of persistence of CD45RA^int^RO^int^ T_SCM_ on CD4+ T cell helper function, longitudinal samples were stimulated with HIV gag peptide pools (**Fig. 5g,h**). Increased frequency of polyfunctional HIV-specific CD4+ T cells (secreting high levels of TNFα and IFNγ) were detected at years 4 and 5 post infusion. Similarly, increased HIV-specific CD8^+^ T cell responses were also detected at years 4 and 5 post infusion (**Fig. 5i,j**). Responding CD4^+^ T cells included cells of T_EM_ phenotype as well as CD45RA^int^RO^int^ CD27^−^CCR7^−^ cells, while responding CD8^+^ T cells included cells of T_TM_, T_EM_, and T_EMRA_ phenotypes (**Supplementary Fig. 9**). Concomitantly to increased HIV-specific T cell responses, a decrease in activation and exhaustion markers was observed in CD8^+^ T cells at years 4 and 5 post infusion (**Supplementary Fig. 10**). To determine how ongoing viral replication impacted the frequency of intact HIV DNA proviruses, the IPDA assay was performed on longitudinal samples. The frequency of intact provirus in purified CD4^+^ T cells remained low post ATI and increased between years 4 and 5 (from 6 to 127 copies per 10^6^ cells; **Fig. 5k**). The low level of VL at year 5 despite an increase in the frequency of intact provirus suggests a role for enhanced CD8^+^ HIV-specific immunity in maintaining control of viral replication.

**Figure 5.**
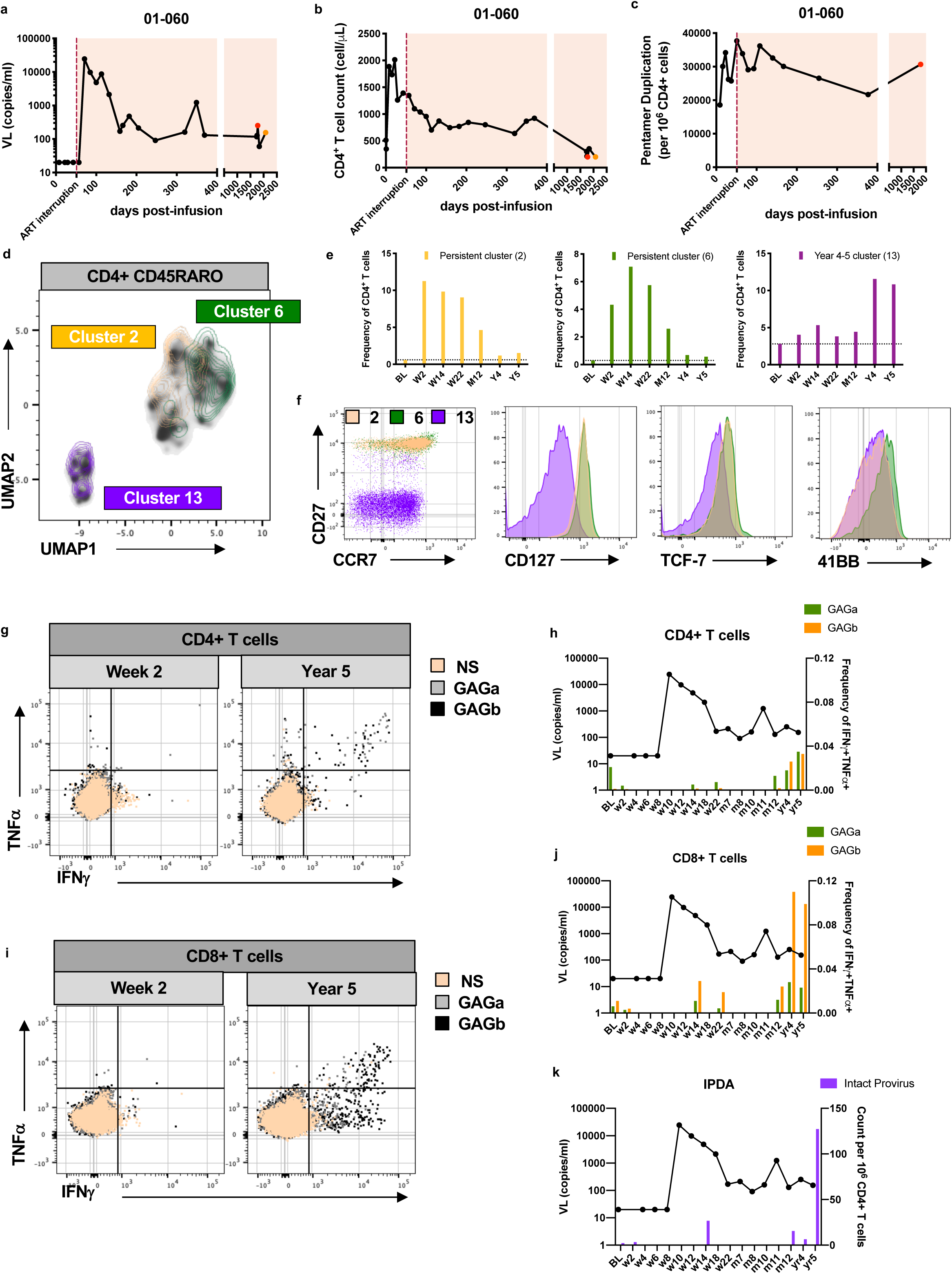
Long term control of viral load for up to 5 years post infusion in participant 01-060. Participant 01-060’s VL (**a**), CD4+ T cell counts (**b**), and Pentamer Duplication levels normalized to CD4+ T cells (**c**) are shown post-infusion, post-ATI, and during long term follow up. The years 4 and 5 time point visits are highlighted in red and orange, respectively. **d,** Uniform Manifold Approximation and Projection (UMAP) dimension reduction analysis of flow cytometry phenotypic analysis of the CD45RA^int^RO^int^ T_SCM_ subset in participant 01-060, with overlays of clusters 2 and 6 (which increased by week 2 post infusion and remain above BL) and 13 (which increased at years 4 and 5) **e**, Frequency of clusters 2, 6 and 13 in total CD4+ T cells. **f**, Expression of markers upregulated in memory stem cells (CD27, CCR7, CD127, TCF-6, and 41BB) in clusters 2, 6 and 13. **g**, Dot plot of CD4+ T cells producing both IFN-γ and TNF-α cytokines post a 6-hour gag peptide pool stimulation in longitudinal samples from participant 01-060 (shown for week 2 and year 5). **h**, The frequency of CD4^+^ T cells producing both IFN-γ and TNF-α cytokines post a 6-hour gag peptide pool stimulation in longitudinal samples from participant 01-060 is overlaid with VL levels post ATI. **i**, Dot plot of CD8^+^ T cells producing both IFN-γ and TNF-α cytokines post a 6-hour gag peptide pool stimulation in longitudinal samples from participant 01-060 (shown for week 2 and year 5). **j**, The frequency of CD8+ T cells producing both IFN-γ and TNF-α cytokines post a 6-hour gag peptide (GAGa ad GAGb) pool stimulation (green and orange bars, respectively; right Y axis) in longitudinal samples from participant 01-060 is overlaid with VL levels post ATI. **k**, Frequency of intact provirus as measured by the IPDA assay (purple bars; right Y axis) in longitudinal samples from participant 01-060 is overlaid with VL levels (black).

## Discussion

Previous studies of non-modified autologous T cell infusion in HIV-infected humans did not result in sustained CD4^+^ T cell reconstitution^28,43,44^ nor HIV reservoir decay, partly due to minimal persistence of infused cells^45^ and their susceptibility to infection^46^. Herein we show in two independent clinical trials that a single infusion of autologous CCR5 gene-edited cells is safe and well tolerated (see online discussion). Importantly, we found in the first study (SB-728-0902) that this intervention led to a sustained albeit quantitively modest increase in CD4^+^ T cell numbers in individuals who failed to normalize their CD4^+^ T cell counts despite long-term effective ART, as well as to a significant long-term decay of the estimated size of the total HIV reservoir, with a decrease in HIV DNA of over 1 log_10_ copies per million cells in 4 of the 9 individuals. The decrease in HIV-infected cells using measures of HIV DNA is in sharp contrast to the very stable levels reported during long-term ART^9,10,47^ and in recent clinical trials using latency reversal agents^48–52^ and was not a result of dilution of HIV-infected cells by the expansion of the infused product as the majority of the HIV decay was continuous over a span of 3 years. Our observed outcomes were associated with the expansion of a novel T_SCM_ subset, CD45RA^int^RO^int^ T_SCM_, that persisted long-term and comprised between 3 and 26% of CD4^+^ T cells 3-4 years post-infusion. The association of the expansion of this novel CD45RA^int^RO^int^ T_SCM_ subset with improved control of HIV replication following ART interruption was demonstrated in the SB-728-1101 study. Our results highlighted the capacity of CD45RA^int^RO^int^ T_SCM_ to differentiate into downstream short-lived memory T cells, by tracking ZFN-mediated mutations specific to CD45RA^int^RO^int^ T_SCM_, as well as the enhanced HIV-specific CD8^+^ T cell responses through cognate or non-cognate help as mechanisms associated with reservoir decay and virological control.

A CD45RA^+^CD45RO^−^CCR7^+^CD27^+^CD95^+^ T_SCM_-like phenotype was reported following *in vitro* expansion of purified CD4^+^ and CD8^+^ naive T cells^35,53^ and *in vivo*^31^. This CD45RA^+^ T_SCM_ subset was shown to be endowed with self-renewal properties in serial transplantation experiments and long-term persistence (reviewed by Gattinoni et al^54^). The CD45RA^int^RO^int^ T_SCM_ subset described in both our clinical studies was present in the SB-728-T products and proliferated post-infusion as these cells significantly increased in cell frequencies. We demonstrated the importance of the longevity of CD45RA^int^RO^int^ T_SCM_ in the long-term persistence of infused CCR5 gene-edited cells. The capacity of CD45RA^int^RO^int^ T_SCM_ for self-renewal was indicated by the presence of identical CCR5 mutations in the products and 3-4 years post-infusion and the maintenance of a significant fraction (∼25%) of gene-edited cells in this subset 3-4 years post-infusion. The importance of this stem cell phenotype for increased CD4^+^ T cell numbers and HIV reservoir decay was confirmed as this subset was positively associated with both clinical outcomes, as well as with VL control.

Tracking of CCR5 ZFN-mediated mutations showed that CD45RA^int^RO^int^ T_SCM_ cells have the capacity to differentiate into other memory subsets including T_EM_ (which included up to 3% of gene-edited alleles at years 3-4 post-infusion) defining this subset as a bona fide multipotent memory stem cell. Similar monitoring approaches of gene edited cells, using retroviral integration site (IS) analysis^44,55–58^, have shown the differentiation of HSC to naive T cells and demonstrated the transition of CD45RA^+^ T_SCM_ to T_CM_ post-infusion of gene-edited cells. The potential of CD45RA^int^RO^int^ T_SCM_ to form a diverse progeny was confirmed *in vitro* following TCR stimulation of this subset and phenotypic evaluation of proliferating cells. A multivariate model analysis of predictors of the HIV reservoir decline confirmed that long-term persistence of CD45RA^int^RO^int^ T_SCM_ and differentiation of CCR5 gene-edited T_SCM_ into T_EM_ were associated with the reduction of HIV-infected cells. Our results suggest that the long-term decrease in HIV DNA post-infusion is a consequence of the continuous replacement of the pool of short lived T_EM_ cells, known to be preferentially infected by R5 tropic viruses^59^, by the progeny of T_SCM_ cells that have low frequencies of HIV-infected cells and hence would not replenish the viral reservoir; moreover, these cells harbor CCR5 gene-edited alleles which also makes their progeny resistant to infection.

CD45RA^+^ T_SCM_ have been shown to be susceptible to *in vitro* infection^60^ and can constitute a stable source of latent HIV reservoir^61,62^. A central hypothesis of our studies was that providing protection from HIV infection to a subset of CD4^+^ T cells would provide a global benefit and allow for reduction of the latent HIV reservoir and control of viral replication.

We recognized that the decay in HIV DNA in the SB-728-0902 trial might be due to the *ex vivo* expansion of both CCR5 gene edited and gene unmodified T_SCM_ cells within the infused products and that such cells would be protected *in vivo* by ART. Although several reports have described active viral replication during suppressive ART^63–65^, this remains a controversial subject^66,67^. Nonetheless, inefficient drug penetrance, impaired immune function, immune privileged sites, and release of virus through clonal proliferation of infected cells^16,68^ have been suggested as mechanisms for viral persistence^69,70^. Consequently, CCR5 gene editing would offer a selective advantage in the presence of ongoing viral production in potential sanctuary sites. Nevertheless, our results from the SB-728-1101 trial demonstrate an increase in the frequency of CCR5 gene-edited cells in T_EM_ and a lack of an increase in integrated HIV DNA in that subset during active viral replication in PTCs, highlighting that differentiation of CCR5 gene-edited T_SCM_ into other memory subsets can lead to protection from *de novo* HIV infection. This was further confirmed by long-term monitoring of the pentamer duplication marker in the absence of ART in participant 01-060 where we saw an enrichment of CCR5 gene-edited cells between years 1 and 4 post-infusion. The increase in intact proviral DNA at year 5 confirms ongoing viral replication. Interestingly, this individual was heterozygote for the delta-32 CCR5 gene mutation which may have resulted in a greater probability of circulating bi-allelic CCR5 deleted cells, which would represent fully protected cells. The importance of limiting HIV infection in T_SCM_ (or T_CM_) cells for the preservation of CD4^+^ T cell homeostasis was shown in non-human primates^71^ and viremic non-progressor HIV-infected individuals^72^. A follow-up randomized clinical trial (NCT03666871) with a larger cohort and a control arm will further evaluate the impact of CCR5 gene modification on T cell homeostasis and reservoir decay in the absence of ATI.

HIV-specific CD8^+^ T cell function has been implicated in the control of viral replication in elite controllers^73–75^ as well as post-ART controllers^76–78^. Moreover, numerous studies have shown that CD4^+^ T cell help is crucial for CD8^+^ T cell function^79,80^. Monitoring T cell responses in participant 01-060 who controlled VL to levels ∼100 copies/ml showed that HIV-specific CD4^+^ T cells producing TNF-a and IFN-g were increased during ongoing ATI and coincided with an increase in HIV-specific CD8^+^ T cell polyfunctional responses. These results are in line with a recent study demonstrating that memory stem cell generation following vaccination in melanoma patients was associated with robust anti-tumor cell responses^81^. We also show in the SB-728-1101 study that HIV-specific CD8^+^ T cell polyfunctional responses increased post-ATI and correlated with HIV viral load and levels of integrated HIV DNA in T_EM_ cells. As only a subset of T_CM_, T_TM_, and T_EM_ cells contain CCR5 mutations, enhanced HIV-specific CD8^+^ T cell responses post-infusion is a mechanism that controls and potentially reduces the HIV reservoir during active viral replication.

In summary, our results indicate that infusion of CCR5 gene-edited cells provides a unique therapeutic intervention that improves T cell homeostasis and reduces the HIV reservoir. The less-invasive and autologous aspect of this therapy makes it more accessible than hematopoietic stem cell transplantation. Combining this approach with other interventions that enhance T_SCM_ proliferation and differentiation post-infusion might further improve outcomes.

## Acknowledgements

We thank the study participants for their participation in this clinical trial. We thank <u>Filipa Blasco Tavares,</u> Vinicius Suzart, <u>Victor Joo,</u> Yuwei Zhang, Franck Dupuy, and Alessandra Noto for technical assistance, Michael Cartwright, Navnita Dutta, and Petra Stafova for assistance with Illumina microarrays, as well as Malika Aid, <u>Abdelali Filali-Mouhim,</u> Courtney Steel and Peter Wilkinson for bioinformatics support. We are grateful to Zhong He, Yu Shi, and Kim Kusser for flow cytometric cell sorting, and to Benjamin A. Youngblood, Mirko Paiardini and Joseph M. McCune for comments on the manuscript. The work was supported by the Foundation for AIDS Research (amfAR grants # 108549-53-RGRL and 108833-55-RGRL) and the Delaney AIDS Research Enterprise to cure HIV (DARE grant # AI096109).

## Author contributions

J.L. recruited participants into the study. D.A. oversaw the clinical trial, clinical data management and clinical operations. S.D. wrote the protocol and oversaw the clinical study with D.A. J.Z., A.S., and G.L. designed and performed experiments and analyzed data under the guidance of R.-P.S. and D.A. A.R. performed statistical analysis of HIV reservoir decay under the guidance of G.M. and J.H. R.F., F.A.P., and N.C. designed and performed HIV reservoir measurements. S.F. performed bioinformatic analysis of CCR5 sequence tracking as well as multivariate model analysis. K.G., A.S., and M.A. performed bioinformatic analysis of gene array results. R.B. provided antibodies and helped with selection of Ab/fluorochome combinations for multiparametric flow panels. G.P.S., G.C., and C.B. contributed to flow cytometry experiments and cell sorting. J.Z. and A.S. prepared the figures. J.Z., L.S., R.-P.S. and S.D. wrote the manuscript.

## Author information

Microarray data was deposited at the GEO database (http://www.ncbi.nlm.nih.gov/geo/), accession number GSE66214. Reprints and permissions information is available at www.nature.com/reprints. The authors declare no competing financial interests. Authors affiliated with Sangamo Therapeutics are employees and investigators are paid by Sangamo to perform the trial according to FDA approved good clinical trial practices. Correspondence and requests for materials should be addressed to R.-P.S.

## Methods

### Study design

The SB-728-0902 clinical trial is a Phase 1, open label, uncontrolled, nonrandomized study of individuals with chronic HIV infection treated with ART (ClinicalTrials.gov # NCT01044654). The study was sponsored by Sangamo Therapeutics and was conducted at two centers in the United States between December 2009 and April 2014. The primary objective of the study was to assess the safety and tolerability of ascending dose of autologous CD4^+^ enriched T cells edited at the CCR5 gene by ZFNs (SB-728-T cells). Secondary objectives included the assessment of increases in CD4^+^ T cell counts, long-term persistence of CCR5 gene-edited cells, homing to gut mucosa, and the effects on HIV viral persistence (HIV RNA and proviral DNA). A total of 9 participants were enrolled into three ascending dose cohorts (Cohort 1 received 10 x 10^9^ SB-728-T, cohort 2 received 20 x 10^9^ SB-728-T, and cohort 3 received 20 x 10^9^ SB-728-T), with three participants in each cohort (Supplementary Table 1). All participants were followed weekly for the initial 4 weeks and then monthly thereafter for one year, after which they were enrolled in a three-year safety study. Participant 1-01 underwent a treatment interruption between months 12 and 22; viral load measurements are listed in Supplementary Tables 10 and 11 and ART regimens are detailed in Supplementary Table 13.

The SB-728-1101 clinical trial is a Phase 1, open label, uncontrolled, nonrandomized study of individuals with chronic HIV infection treated with ART (ClinicalTrials.gov #NCT01543152). The study was sponsored by Sangamo Therapeutics and was conducted at 12 centers in the United States between March 2012 and January 2017. The primary objective of the study was to evaluate the safety and tolerability of escalating doses of cyclophosphamide (CTX) pre-treatment to promote CD4^+^ T cell expansion after administration of a single dose of SB-728-T cells. Participants received CTX at doses of 0.1 (Cohort 1, *n* = 2), 0.5 (Cohort 2, *n* = 4), 1.0 (Cohort 3, *n* = 3), 1.5 (Cohort 5, *n* = 3) and 2.0 g/m^2^ (Cohort 4, *n* = 3) one day before infusion of SB-728-T cells (Supplementary Table 14). Participants subsequently received between ∼10 to 40 billion SB-728-T cells (Supplementary Table 8). All participants were followed weekly for the initial 4 weeks, bi-monthly until week 14, monthly until week 22, and then every 2 months until month 12. ART was discontinued 6 weeks after SB-728-T infusion for a period of 16 weeks (Supplementary Fig. 7b). Secondary objectives included the evaluation of the effect of SB-728-T cells on plasma HIV-1 RNA levels following ART interruption. During the treatment interruption, ART was reinstituted in participants whose CD4^+^ T cell counts dropped to <500 cells/µL and/or whose HIV-RNA increased to >100,000 copies/mL on three consecutive weekly measurements. Following completion of the 1-year study, participants were enrolled in a 3-year long-term safety study. Adverse events are summarized in the on-line discussion.

The final clinical protocol, amendments, and consent documents were reviewed and approved by the NIH Recombinant DNA Advisory committee, as well as institutional review board and institutional biosafety committee (as required) at each study center. All participants provided written informed consent.

### Enrollment criteria

#### SB-728-0902 Trial

Eligible participants were 18 years of age or older and were chronically infected with HIV, as documented by ELISA. Participants were on long-term stable ART and aviremic (undetectable HIV RNA for at least one year prior to enrollment), with CD4^+^ T cell counts between 200 and 500 cell/µL (immune non-responders; Supplementary Table 1) and had adequate venous access and no contraindications to leukapheresis. The key exclusion criteria included a SNP at the CCR5 ZFN target region, current or prior AIDS diagnosis, receiving therapy with maraviroc or immunosuppressives, and hepatitis B or hepatitis C co-infection.

#### SB-728-1101 Trial

Eligible participants were 18 years of age or older and were chronically infected with HIV, as documented by ELISA. Participants were aviremic on stable ART with CD4^+^ T cell counts >500/µL, had R5 tropic HIV, and willing to discontinue current ART during the treatment interruption (Supplementary Table 14). The key exclusion criteria included adenoviral neutralizing antibodies >40, a SNP at the CCR5 ZFN target region, current or prior AIDS diagnosis, receiving therapy with maraviroc or immunosuppressives, and hepatitis B or hepatitis C co-infection.

### Cell manufacture and infusion

Production of SB-728-T cells was previously described^29,82^. Briefly, participants underwent a 10L leukapheresis to collect, enrich, modify and expand autologous CD4^+^ T cells. Manufacturing of the SB-728-T products includes a T cell activation step (with anti-CD3/anti-CD28-coated magnetic beads) and an IL-2 expansion step^29^. ARV drugs (Ritanovir and Norvir with or without Fuzeon) were added during manufacture to inhibit de novo infection of CD4^+^ T cells. SB-728-T refers to autologous CD4^+^ enriched T cells that have been transduced *ex vivo* with SB-728, a replication deficient recombinant Ad5/35 viral vector encoding the CCR5 specific ZFNs (SBS8196z and SBS8267), that includes a mixture of gene-edited and non-edited cells. Expression of CCR5-specific ZFNs induces a double stranded break in the cell’s DNA which is repaired by cellular machinery leading to random sequence insertions or deletions (indels) in ∼25% of transduced cells. These indels disrupt the CCR5 coding sequence leading to frameshift mutation and termination of protein expression.

Infusion of SB-728-T was conducted employing a standard intravenous infusion method common to all adoptive cell transfer protocols. Participants were pre-medicated with acetaminophen 650 mg P.O. and 25-50 mg Benadryl P.O. approximately 1 hour prior to infusion. Cryopreserved SB-728-T products were thawed bed-side and infused via the intravenous route using the Smartsite gravity set.

### Cryopreserved peripheral blood mononuclear cells (PBMCs) samples

#### SB-728-0902

PBMCs were prepared from whole blood by ficoll-hypaque density sedimentation and used cryopreserved in 10% dimethyl sulfoxide (DMSO) and 90% FBS. Availability of cryopreserved samples at different time points varied between participants; consequently, time points were grouped into early (14-28 days), mid (4-7 months or 9-10 months), late (11-12 months), and long-term (2-3 or 3-4 years) post-infusion time points. Baseline samples included cryopreserved PBMCs from the initial leukapheresis (2-3 months before infusion) as well as from a small volume blood draw 1-weeks before infusion. PBMCs from participants 1-01, 1-02, and 1-03 were not cryopreserved until months 6 or 8 post-infusion. Most participants agreed to a large volume blood draw (*n* = 9, year 2-3) and a leukapheresis (*n* = 7, year 3-4) during the long-term follow-up period to allow for assays requiring large amounts of cells, such as CCR5 sequencing, integrated HIV DNA quantification, and gene arrays in sorted CD4^+^ T cell subsets. For certain assays, including flow cytometry phenotyping, baseline samples for only 6 participants remained available. Manufacturing samples (SB-728-T products) were also available for all participants.

#### SB-728-1101

Clinical measures (CD4, CD8 counts, viral load (VL), and the Pentamer Duplication marker) were performed at every time point. Availability of cryopreserved PBMCs at baseline and pre-ATI were not available for Cohorts 1 and 2 participants; consequently immunological (T cell phenotyping, CCR5 DNA sequencing of ZFN mediated mutations in sorted CD4^+^ subsets) and virological (Integrated HIV DNA) measurements were only performed in participants from Cohorts 3-5. Baseline samples included cryopreserved PBMCs from the initial leukapheresis (2-3 months before infusion) as well as from a small volume blood drawn 1-2 weeks before infusion. Manufacturing samples (SB-728-T products) were also available for Cohorts 3-5 participants.

### Rectal and lymph node biopsies

Rectal biopsies were performed for participants of the SB-928-0902 trial at baseline, day 14, month 3, 6 and 12 (*n* varied between 3 and 9 participants per time point). Mucosal mononuclear cells were isolated from sigmoid colon biopsies obtained by endoscopy via a combination of collagenase digestion and teasing with 18G needles. Inguinal lymph nodes were biopsied from 3 volunteers at one time point (between 9 and 18 months) post-SB-728-T infusion. Tissues were processed into single cells as described in Anton et al^83^ and genomic DNA were isolated for assessment of CCR5 gene modifications.

### Quantification of *CCR5* gene modification in SB-728-T products using Cel-I

The Cel-I nuclease specifically cleaves DNA duplexes at the sites of distortions created by either bulges or mismatches in the double helical DNA structure. We have adapted protocols using this enzyme for quantification of minor indels typically induced by ZFN-mediated gene modifications. Briefly, the genomic region of interest (CCR5) is PCR amplified, the PCR product is denatured and then allowed to re-anneal to permit wild type and non-homologous end joining-edited alleles to re-anneal together and create hetero-duplexes. The re-annealed PCR products are then digested with the Cel-I nuclease to cut the PCR-amplified DNA at the site of mismatches. Subsequently, the level of ZFN-mediated gene modification can be quantified by determining the ratio of the uncleaved parental fragment to the two lower migrating cleaved products.

### Quantification of CCR5 gene-edited CD4^+^ T cells by Polymerase Chain Reaction

ZFN-mediated gene modification can generate a wide range of frame-shift mutations to disrupt the CCR5 gene locus. A PCR-based assay was developed to measure the acquisition of a unique duplication of 5-nucleotide (Pentamer) DNA sequence, CTGAT, at the ZFN cleavage site in approximately 25% of the gene-edited alleles^29^. Genomic DNA (gDNA) was extracted from PBMCs using a commercially available kit (Masterpure DNA Purification kit, Epicenter, Madison, WI). A standard PCR was performed with 5µg of gDNA to amplify a 1.1 kb region that contains CCR5 gene modifications. This 1.1 kb amplicon is subsequently evaluated with the two independent qPCRs, one specific for the Pentamer Duplication-CCR5 gene-edited allele (by using a primer that contains the Pentamer Duplication), and a second that amplifies all CCR5 alleles. The ratio of Pentamer Duplication-specific templates and the total number of CCR5 alleles yields Pentamer Duplications per 1 million PBMCs. The assay has a sensitivity of one CCR5 gene-edited allele per 10^5^ total CCR5 alleles. As the Pentamer Duplication markers represents ∼25% of ZFN-mediated gene modifications, the total frequency of CCR5 gene-edited cells in PBMCs was estimated by multiplying the frequency of Pentamer Duplication gene-edited cells by 4.

### Estimation of expansion of SB-728-T post-infusion

The level of CCR5 gene-edited alleles present in a participant post-infusion relative to the amount infused can be estimated using the measured values of CCR5 modification by the Pentamer Duplication marker and CD4^+^ cell count with the assumptions; 1) blood volume is 4.7 liters, 2) approximately 2.5% of all CD4^+^ T cells are found in the periphery^84^ and 3) SB-728-T products distribution is similar to endogenous CD4^+^ T cells (levels of CCR5 modification in CD4^+^ T cells from the sigmoid and inguinal nodes are similar to that in the periphery, Supplementary Fig. 4).

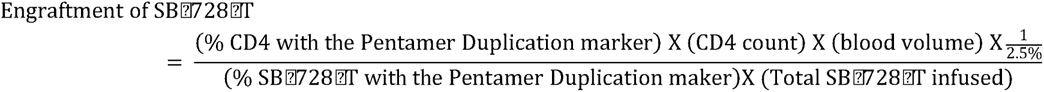

### Quantification of *CCR5* gene modification via next-generation sequencing/MiSeq

The locus of interest (ZFN binding sites in *CCR5*) was PCR amplified from genomic DNA, and the levels of modification at each locus were determined by paired-end deep sequencing on an Illumina MiSeq sequencer. Paired sequences were merged via SeqPrep (John St. John, https://github.com/jstjohn/SeqPrep, unpublished). A Needleman-Wunsch alignment was performed between the target amplicon genomic region and the obtained Illumina sequence to map indels^85^. CCR5 sequencing was performed in sorted CD4^+^ T cell subsets from SB-728-0902 participants in SB-728-T products (*n* = 9) and year 3-4 samples (*n* = 8) as well as from SB-728-1101 Cohorts 3-5 participants in SB-728-T products (*n* = 7) and weeks 6 and 22 samples (*n* = 7). Perez *et al.,* had shown that approximately 30% of CCR5 gene modifications were bi-allelic^29^. For simplicity, CCR5 gene-edited memory subset cell counts were estimated by multiplying each memory subset cell count by the frequency of CCR5 gene-edited alleles within each memory subset as determined by CCR5 DNA-Seq, thereby assuming that one gene-edited allele is equivalent to one cell.

### Cell tracking of CCR5 gene-edited CD45RA^int^RO^int^ T_SCM_ post SB-728-T infusion

Sequencing of CCR5 ZFN-mediated mutations in sorted CD4^+^ T cell subsets was used to track differentiation of CD45RA^int^RO^int^ T_SCM_ cells post-infusion. First, wild type CCR5 (amplicon) and sequences detected in only 1 of the samples sequenced were excluded from further analysis (∼80% of unique CCR5 sequences). Then, for each participant, we identified the sequences expressed only in CD45RA^int^RO^int^ T_SCM_ cells in SB-728-T products and then analyzed their distribution in CD4^+^ T cell memory subsets at year 3-4 samples (SB-728-0902) or weeks 6 and 22 samples (SB-728-1101).

### Flow cytometry analysis

#### Surface staining

Two surface panels were run for longitudinal analysis of CD4^+^ T cell distribution in SB-728-T products and post-infusion, including a T_SCM_ panel (Figs. 1a,c and 4c, and Supplementary Fig. 3), using previously titrated monoclonal antibodies summarized in Supplementary Table 8. Thawed PBMCs (1-2 million cells) were labelled for 30 minutes in the dark at 4°C, washed with staining buffer (PBS, 2% FBS), fixed with 2% FA (Sigma Aldrich) for 15 min at 22°C, and resuspended in staining buffer for acquisition. A minimum of 100,000 live cells were acquired within 24hrs using a BD LSR-II or BD LSRFortessa™ cell analyzer and analyzed using the FlowJo version 9 software (TreeStar, Ashland, OR). Longitudinal samples from each participant were stained in the same batch run and a common set of control cells (obtained from an independent ART-treated HIV-infected donor) was stained when samples from different participants were stained in separate runs.

#### Intracellular Cytokine Staining (ICS)

A transcription factor panel (Supplementary Table 8) was used to determine Th-lineage in CD4^+^ T cell subsets in the SB-728-0902 study (Fig. 3c). Thawed PBMCs were labelled with surface antibodies for 30 minutes at 4°C prior to fixation with eBioscience’s Foxp3 Fixation/Permeabilization buffer for 30 minutes at 4°C. Cells were then labelled with the intracellular antibodies for 30 minutes at 22°C in eBioscience’s Foxp3 permeabilization buffer, resuspended in staining buffer (PBS, 2% FBS), and immediately acquired using a BD LSRFortessa™ flow cytometer.

An ICS panel (Supplementary Table 8) was used to quantify effector cytokine production in CD4^+^ T cell subsets following T cell stimulation (Fig. 3d-g) or gag peptide pool stimulation (Fig. 4g-j and Supplementary Fig. 9). Cells were labelled with surface antibodies, and permeabilised with Perm buffer (BD Bioscience) after which cells were stained intracellularly prior to fixation with 2% formaldehyde. Cells were acquired within 24 hours using a BD LSR-II or BD LSRFortessa™. A minimum of 500,000 live events was acquired. Longitudinal samples were stained in the same batch run. Cells were analyzed using FlowJo version 10.

Additional intracellular panels were used to further characterize the stemness of the CD45RA^int^RO^int^ subset and CD8^+^ T cell activation and exhaustion (Fig 4d-f and Supplementary Fig. 10) in longitudinal samples from participant 01-060 (Supplementary Table 8).

Cell proliferation and stemness/effector phenotype progression of *in vitro* stimulated CD45RA^int^RO^int^ T_SCM_ (Fig. 1e-h and Supplementary Fig. 4) was determined by measuring the progressive dilution of CellTrace Violet (Thermo Fisher Scientific) combined with an intracellular panel (Supplementary Table 8).

### Uniform Manifold Approximation and Projection (UMAP) Analysis of flow cytometry panels

Live CD3^+^CD4^+^ cells (Figs. 1e-h, 3d-g, and Supplementary Fig. 4), live CD4^+^CD45RA^int^RO^int^ cells (Fig. 5d-f and Supplementary Fig. 8c-e), or live CD3^+^CD4^−^CD8^+^ T cells (Supplementary Fig. 10) were gated and exported for unbiased clustering analyses. For these panels, projection of the density of cells expressing markers of interest were visualized/plotted on a 2-dimensional UMAP (https://arxiv.org/abs/1802.03426, https://github.com/lmcinnes/umap). Clusters of cells were identified using the RPhenograph package (https://github.com/jacoblevine/PhenoGraph) after concatenating all samples per panel and bi-exponentially transforming each marker. The K value, indicating the number of nearest neighbors, was set to 60. Data were visualized using FlowJo version 10 and R for heatmaps highlighting differences in MFI for each marker per cluster.

### Cell culture and stimulation conditions

Thawed PBMCs were rested in RPMI 1640 medium supplemented with 10% FBS and 1% penicillin–streptomycin for 12 hours. To induce cytokine production, 2 million cells were activated with dynabeads human T activator CD3/CD28 at a 1:1 bead to cell ratio (Thermo Fisher Scientific), Staphylococcal enterotoxin B (SEB; 1µg/mL) (Toxin Technology), Phorbol myristate acetate (PMA) (100 ng/mL) and Ionomycin (1µg/mL) (both from Sigma Aldrich), or only complete media (mock) for 6 hours in the presence of Brefeldin A (5µg/mL) (Sigma Aldrich). In a separate experiment, thawed PBMCs were rested for 12 hours prior to stimulation of 2 million cells with either gag peptide pool (GAGa and b pools; 1µg/mL; NIH AIDS reagent program), Staphylococcal enterotoxin B (SEB; 1µg/mL) (Toxin Technology), CEF control peptide pool (SB-728-1101 study only: 1µg/mL; NIH AIDS reagent program), or only complete media (mock) for 6 hours in the presence of Brefeldin A (5µg/mL) (Sigma Aldrich).

### HIV DNA in PBMCs, total and sorted CD4^+^ T cell subsets

Total HIV DNA in PBMCs was measured by droplet digital polymerase chain reaction. In brief, genomic DNA (gDNA) was extracted from PBMCs using a commercially available kit (Masterpure DNA Purification kit, Epicenter, Madison, WI). 2 µg of gDNA was digested with the restriction enzyme DdeI at 37°C for 1 hour. PCR droplets were prepared according to manufacturer’s recommendations. Briefly, a 20µL of multiplex PCR mixture is prepared by mixing 250 or 500 ng of the digested gDNA with the ddPCR™ 2x Master Mix and two Taqman primer/probe sets. The Taqman primer/probe sets amplify a conserved region in *gag* (as described by Palmer *et. al.*, HIV gag forward CATGTTTTCAGCATTATCAGAAGGA, HIV gag reverse TGCTTGATGTCCCCCCACT, HIV gag probe, FAM-CCACCCCA CAAGATTTAAACACCATGCTAA-BHQ) and the human Ribonuclease P protein subunit p30 (RPP30 forward GATTTGGACCTGCGAGCG, RPP30 reverse GCGGCTGTCTCCACAAGT, RPP30 probe VIC-CTGACCTGAAGGCTCT-MGB-BHQ)^86^. PCR droplets were generated in a DG8™cartridge using the QX-100 droplet generator, where each 20µL PCR mixture was partitioned into approximately 15,000 nano-liter size droplets. PCR droplets were transferred into a 96-well PCR plate and sealed with foil. Standard PCR was performed with a Bio-Rad C1000 Thermal Cycler (95°C (60sec), 40 cycles of 94°C (30sec)/ 60°C (60sec), 98°C (600 sec)). HIV DNA copy number was evaluated using the QX-100 Droplet Digital PCR system (Bio-Rad, Hercules, CA). The PCR-positive and PCR-negative droplets for HIV gag and RPP30 were determined and template concentrations were calculated by Poisson analysis. HIV copy number was determined by normalizing HIV gag concentration to RPP30 concentration^86^. Integrated DNA was measured as previously described^87^ in purified CD4^+^ T cells from SB-728-0902 participants at baseline, in SB-728-T products, year 2-3 samples (*n* = 9) and in sorted CD4^+^ T cell subsets in SB-728-T products and year 3-4 samples (*n* = 8). Integrated DNA was also measured in purified CD4^+^ T cells as well as sorted CD4^+^ T cell subsets from SB-728-1101 Cohorts 3-5 participants in baseline, weeks 2-6 and 14-22 samples (*n* = 8).

### HIV Tropism Assay

HIV Tropism was evaluated using the commercial Trofile® DNA assay (Monogram BioSciences/ LabCorp, South San Francisco, CA). Viral envelope DNA sequence was extracted from PBMCs. HIV tropism is determined using a cell-based transduction assay where HIV env protein sequences are amplified from PBMC samples, subcloned as a library, packaged into lentiviral vectors, and evaluated using co-receptor restricted cell lines.

### T cell Receptor (TCR) Repertoire

TCR repertoire analysis was performed with the immunoSEQ assay (Adaptive Biotechnologies, Seattle, WA). The immunoSEQ method amplifies rearranged TCR CDR3 sequences by multiplex PCR to explore all Vβ and Jβ combinations from isolated genomic DNA and uses high-throughput sequencing technology to sequence TCR CDR3 chains to determine the composition of various T cell clones within each sample. TCR diversity is assessed using the Shannon entropy index^33,88^, which accounts for both the number of unique clones (richness) and clone distribution (evenness) of the TCR Vβ CDR3 sequences present in each sample. A larger Shannon entropy index reflects a more diverse distribution of the TCR Vβ CDR3 sequences.

### Cell sorting

For quantification of the Pentamer Duplication maker, CCR5 gene-edited alleles using DNA-Seq, and levels of integrated HIV DNA within CD4^+^ T cell subsets, CD4^+^ T cells were first isolated from PBMCs by negative magnetic selection (StemCell), and then surface stained with CD3 Alexa 700 (clone UCHT1), CD95 PE-Cy7 (clone DX2), CD58 PE (clone 1C3), CD127 BV421 (clone HIL-7R-M21), CD28 APC (clone CD28.2), CD14 V500 (clone M5E2) (all from BD Biosciences), CD4 Qdot 605 (clone S3.5) (Invitrogen), CD27 APCe780 (clone O323) (eBioscience), CD45RA BV 650 (clone HI100), CD45RO PerCPe710 (clone UCHL1), CD19 BV 510 (clone H1B19) (all from Biolegend), CCR7 FITC (clone 150503) (R&D), aqua fluorescent reactive dye (Invitrogen), and CD8 PerCP (clone SK1) from Biologend (SB-728-0902 study) or CD8 BV711 (clone RPA-T8) from BD (SB-728-1101 study). Up to 200,000 total CD4^+^ T cells as well as CD4^+^ T cell subsets were then sorted with the FACSAria (Becton Dickinson) and stored as dry pellets at −80°C until analysis. For gene array analysis of CD4^+^ T cell subsets in the SB-728-0902 study, 10,000 sorted cells (naïve, CD45RA^+^ T_SCM_, CD45RA^int^RO^int^ T_SCM_, T_CM_, and T_EM_) were collected directly into RNAse-free 1.5mL eppendorf tubes containing 500µL of RLT buffer with 1% β-mercaptoethanol and stored at −80°C until analysis.

### Gene Microarray and Analyses

Sorted CD4^+^ memory subsets from year 3-4 samples were sorted into RLT buffer as described above and included naïve, CD45RA^+^ T_SCM_, CD45RA^int^RO^int^ T_SCM_, T_CM_, and T_EM_ cells. Sorted cells were lysed for RNA extraction as per manufacturer’s instructions (Qiagen, Valencia, CA). T7 oligo(dT) primed reverse transcription reactions were performed followed by *in vitro* transcription. These products underwent a second round of amplification (MessageAmp II aRNA Amplification kit by Life Technologies) yielding biotin-labeled aRNAs which were hybridized to the Illumina Human HT-12 version 4 Expression BeadChip according to the manufacturer’s instructions and quantified using an Illumina iScan System.

Analysis of gene array output data was conducted using the R statistical language (http://www.r-project.org/)89 and the Linear Models for Microarray Data (LIMMA) statistical package^90^ from Bioconductor^91^. Briefly, scanned array images were inspected for artifacts and unusual signal distribution within chips, and arrays with low overall intensity or variability were removed from analysis. Diagnostic plots such as density plots, box plots, and heatmaps of between-array distances were used to assess hybridization quality across chips. Intensities were log2 transformed before being normalized using the quantile normalization method^92^. Probes that did not map to annotated RefSeq genes and control probes were removed. The LIMMA package was used to fit a linear model to each probe and perform a moderated Student’s *t*-test to assess the difference in gene expression level between the different subsets. For data mining and functional analyses, genes that satisfied a *P* value < 0.05 were selected. The proportions of false positives were controlled using the Benjamini and Hochberg method^93^. All microarray data have been deposited in GEO under accession number GSE66214.

We performed Gene Set Enrichment Analysis (GSEA)^94^ using MSigDB (http://software.broadinstitute.org/gsea/msigdb/) curated gene sets to identify enriched biological pathways that are modulated in CD45RA^int^RO^int^ T_SCM_ compared to the CD45RA^+^ T_SCM_ subset. GSEA is a statistical method to determine whether members of a particular gene set preferentially occur toward the top or bottom of a ranked-ordered gene list where genes are ranked by the strength of their association with the outcome of interest. More specifically, GSEA calculates an enrichment score (NES) that reflects the degree to which a set of genes is overrepresented among genes differently expressed. The significance of an observed NES is obtained by permutation testing: resorting the gene list to determine how often an observed NES occurs by chance. Leading Edge analysis is performed to examine the particular genes of a gene set contributing the most to the enrichment. We discarded gene sets with a false discovery rate (FDR) > 25% and a nominal *P* value > 0.05.

We used linear regression analysis to identify genes in CD45RA^int^RO^int^ T_SCM_ that correlated with the HIV reservoir decay (Ratio of total HIV DNA (log_10_)/10^6^ PBMC at day 720 over day 0). We fit a linear model (using R language) between each of the genes and the levels of these outcomes as continuous variables and used GSEA to associate a pathway positively or negatively with both of the readouts (Fig. 2e, 3b, and Supplementary Tables 6 and 7).

### Statistical Analysis

#### Clinical data (CD4 counts, CD4:CD8 ratio, HIV DNA), CCR5 modification, and flow cytometry analysis

The paired Wilcoxon rank-sum two-tailed test was used to perform non-parametric donor-paired two-sided analysis to assess the significance of post-infusion changes in total CD4^+^ T cell counts, CD4:CD8 ratio, CD4^+^ T cell subset frequencies and counts, and frequency of total and integrated HIV DNA per 10^6^ cells compared to baseline in the SB-728-0902 study. Missing HIV DNA baseline values for participants 1-01 and 1-02 were estimated by the linear regression model fit intercept. The paired Wilcoxon rank-sum two-tailed test was also used to assess the significance of post-ATI changes in CD4^+^ T cell subset frequencies and counts compared to pre-ATI (week 6) and changes in integrated HIV DNA compared to baseline and pre-ATI (week 6) in the SB-728-1101 study. The Wilcoxon rank-sum two-tailed test was also used to compare the frequency of CCR5 gene-edited alleles, obtained by DNA-Seq, in CD4^+^ T cell memory subsets to that of CD45RA^int^RO^int^ T_SCM_ in SB-728-T products and post-infusion samples in the SB-728-0902 and SB-728-1101 studies, as well as to compare the levels of transcription factors in CD4^+^ T cell memory subsets to those in CD45RA^int^RO^int^ T_SCM_ post-infusion, and the frequency of integrated HIV DNA in CD4^+^ T cell memory subsets to that of CD45RA^int^RO^int^ T_SCM_ in SB-728-T products and post-infusion samples in the SB-728-0902 study. The Wilcoxon rank-sum two-tailed test was also used to compare the diversity of CCR5 gene-edited alleles between SB-738-T products and long-term time points in each CD4^+^ T cell memory subset. The Mann-Whitney two-tailed test was used to perform unpaired non-parametric two-sided comparisons in instances where the number of matched participants varied across time points and contained less than 6 matched pairs at a given time point, such as for the frequencies of CD95^+^ cells post-infusion compared to baseline. A *P* value < 0.05 was considered significant. Statistical analysis was performed using GraphPad Prism v7.0.

The Spearman’s rho (ρ) test was used to perform non-parametric correlation analysis to assess the relationship between various measures (e.g., CCR5 gene-edited cells and CD4^+^ T cell subset frequencies and/or counts) and clinical outcomes, including delta CD4^+^ T cell counts post-infusion (SB-728-0902), change in the size of the reservoir calculated using the ratio of year 2 values over baseline post-infusion (SB-728-0902), HIV-specific CD8^+^ T cell effector function post-infusion (SB-728-1101), and control of viral replication post-ATI (SB-728-1101). We controlled for multiple comparison testing by calculating the false discovery rate (FDR) value using the original FDR method of Benjamini and Hochberg^93^. FDR values are provided in Table 1 and Supplementary Table 3. A *P* value < 0.05 and a *Q* value < 0.25 were considered significant. These statistical analyses were performed using GraphPad Prism v7.0.

#### Multivariate analysis

A sparse linear multivariate regression model was built to identify features that predict the change in the frequency HIV-infected cells using the “reduction of total HIV DNA in PBMCs between year 2 and baseline” as dependent variable and CD45RA^int^RO^int^ T_SCM_ cell counts at year 3-4, frequency of Pentamer Duplication mutations in CD45RA^int^RO^int^ T_SCM_ post-infusion, ratio of the frequency of Pentamer Duplication in CD45RA^int^RO^int^ T_SCM_ over the frequency of Pentamer Duplication in T_EM_ at years 3-4, and the number of shared mutations between CD45RA^int^RO^int^ T_SCM_ and T_EM_ as possible independent variables. Features used, and their values are detailed in Supplementary Table 5. The K-nearest neighbor function implemented in the R package ‘impute’ (v1.54.0) was used to infer missing values. Default options (k=10) were used. The feature included in the multivariate model were selected by using the LASSO technique as implemented in the R package ‘glmnet’. Briefly, the subsets of independent variable minimizing the mean-squared-error was optimized by leave-one-out-cross-validation. The final multivariate model was built on the entire cohort using the features selected using the LASSO technique. Student *t*-test was used to test that the regression coefficient of the independent variables of the final model was statistically different from zero. R-squared value was used to assess the proportion of the dependent variable variance explained by the multivariate model.

